# Salt Inducible Kinase activation and IRE1-dependent intracellular ATP depletion to form Sec bodies in Drosophila cells

**DOI:** 10.1101/2021.01.16.426665

**Authors:** Chujun Zhang, Wessel van Leeuwen, Marloes Blotenburg, Angelica Aguilera-Gomez, Sem Brussee, Rianne Grond, Harm H. Kampinga, Catherine Rabouille

## Abstract

The phase separation of the non-membrane bound Sec bodies occurs in Drosophila S2 cells by coalescence of components of the ER exit sites under the stress of amino-acid starvation. Here we address which signaling pathways cause Sec body formation. We find that two pathways are critical. The first is a SIK dependent pathway induced by salt (NaCl) stress in a necessary and sufficient manner. The second is the activation of IRE1 (one of the key kinases mediating the Unfolded Protein Response) by absence of amino- acids, which partly leads to the depletion of intracellular ATP. However, IRE1 activation is not sufficient to induce Sec body formation and needs to be combined to salt stress. This works pioneers the role of SIK in phase transition and re-enforces the role of IRE1 as a metabolic sensor for the level of circulating amino-acids.

## Introduction

Cell compartmentalization is not only mediated by membrane-bound organelles. It also relies on non-membrane bound biomolecular condensates (the so-called membraneless organelles) that populate the nucleus and the cytoplasm.

The formation of membraneless organelles is shown to occur through phase separation that can be driven by stress (such as ER, oxidative, proteostatic, nutrient), resulting in the formation of stress assemblies (van Leeuwen and Rabouille, 2019). Those are mesoscale coalescence of specific and defined components that phase separate. For instance, stress granules can form upon many different types of stresses often associated with the arrest of translation initiation through the phosphorylation of eIF2 alpha (Aulas et al., 2017) (van Leeuwen and Rabouille, 2019) (Aulas et al., 2018).

On the other hand, nutrient stress leads to the formation of many biocondensates. Most of them are RNA based such as stress granules and P-bodies (van Leeuwen and Rabouille, 2019), but some are not. This is the case for glucose starved yeast where metabolic enzymes foci (Munder et al., 2016) (Petrovska et al., 2014) and proteasome storage granules (Peters et al., 2013) (van Leeuwen and Rabouille, 2019) form, as well as Drosophila S2 cells that form Sec bodies under conditions of amino-acid starvation (Zacharogianni et al., 2014).

Sec bodies are related to the inhibition of protein secretion in the early secretory pathway. The early secretory pathway comprises the Endoplasmic Reticulum (ER) where newly synthesized proteins destined to the plasma membrane and the extracellular medium are synthesized. They exit the ER at the ER exit sites (ERES) to reach the Golgi where they are further processed and dispatched to their correct downstream locations. The ERES are characterized by the concentration of COPII coated vesicles whose formation requires 6 proteins, including Sec12 and Sar1, the inner coat Sec23/Sec24 and the outer coat Sec13/Sec31 (Gomez-Navarro and Miller, 2016). In addition, a larger hydrophilic protein Sec16, has been identified as a key regulator of the ERES organization and COPII vesicle budding (Sprangers and Rabouille, 2015). Many additional lines of evidences support the role of Sec16 in optimizing COPII coated vesicle formation and export from the ER (Farhan et al., 2010) (Joo et al., 2016) (Wilhelmi et al., 2016).

Upon the stress of amino-acid starvation in Krebs Ringer Bicarbonate Buffer (KRB), the ERES of Drosophila S2 cells are remodeled into a large round non-membrane bound phase separated Sec bodies. They are typically observed by immunofluorescence after staining of endogenous Sec16, Sec23) and expressed Sec24-GFP) (Zacharogianni et al., 2014) (**see Figure 1A, A’**). Importantly, Sec bodies are very quickly reversible upon stress relief (addition of growth medium). Last, they appear to protect the components of the ERES from degradation (Zacharogianni et al., 2014) and they help cells to survive under conditions of amino-acid shortage (Zacharogianni et al., 2014) (Aguilera-Gomez et al., 2016).

**Figure 1:**
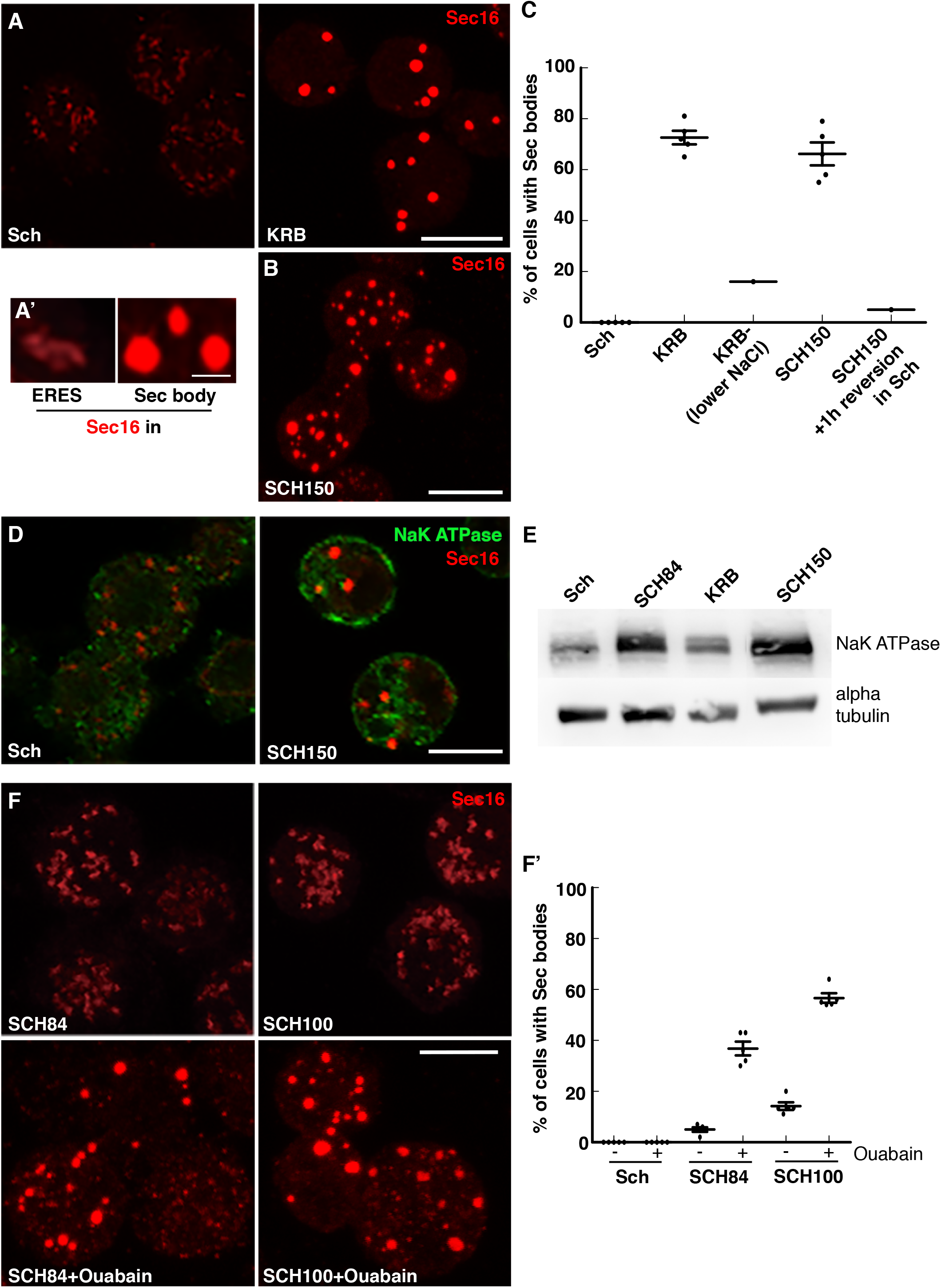
Salt stress activates the Salt Inducible kinases that are involved in Sec body formation. **A, A’**: Immunofluorescence (IF) visualization of endogenous Sec16 (red) in cells growing in Schneider’s (Sch) and in cells incubated in KRB (A). Note the difference in the Sec16 pattern, at ER exit sites in growing cells and in Sec bodies in cells incubated in KRB. There, ERES remodel into larger structures, the Sec bodies, that are brighter than ERES, largely circular and present in at least 2 per cell. Sec bodies are reversible, and EM has previously shown that they are not surrounded by a sealed membrane (Zacharogianni et al., 2014). **B**: IF visualization of Sec body formation (marked by Sec16) in cells incubated in Schneider’s supplemented with 10mM sodium bicarbonate and 150mM of NaCl (SCH150) for 4h at 26°C. **C**: Quantification of Sec body formation (marked by Sec16) in cells incubated in Sch, KRB, KRB-(containing only 60mM NaCl), SCH150 for 4h at 26°C as well as SCH150 reverted in Sch for 1h, showing that SCH150 induced Sec bodies are reversible. **D**: IF visualization of the NaK ATPase (green) and Sec16 (red) in cells incubated in Schneider’s (Sch) and SCH150 for 4h at 26°C. Note the strong increase of NaK ATPase on the cell plasma membrane after incubation in SCH150. **E**: Western blot of S2 cells extract after incubation in Schneider’s (Sch), SCH84, KRB and SCH150 for 4h at 26°C blotted for NaK ATPase and alpha tubulin. Note the NaK ATPase level increased after incubation in SCH150. **F, F’**: IF visualization (F) and quantification (F’) of Sec body formation (marked by Sec16) in cells incubated in Schneider’s, SCH84 and SCH100 supplemented by ouabain (1μM) for 4h at 26°C. Scale bar: 10μm. Errors bars: SEM

Phase separation is shown to be driven by specific components, the so-called drivers, either RNAs and proteins harboring structural features that become “exposed/modified” under certain conditions. In the case of Sec bodies, Sec24A/B (Zacharogianni et al., 2014) (Aguilera-Gomez et al., 2016) and Sec16 have been shown drive Sec body coalescence (Aguilera-Gomez et al., 2016) in a manner that depends on a Sec16 small stretch of 44 residues and on the Mono-ADP-ribosylation enzyme by dPARP16 (Aguilera-Gomez et al., 2016). This illustrates the critical role of post-translational modifications in phase separation (Owen and Shewmaker, 2019) (Bah and Forman-Kay).

In parallel, changes in the cytoplasmic biophysical properties have also been shown to be important in phase separation (Rabouille and Alberti, 2017). For instance, both the formation of metabolic enzyme foci and proteasome storage granules in yeast upon glucose starvation are consequent to a drop of cytoplasmic pH within minute without post-translational modifications (Peters et al., 2013) (Munder et al., 2016) (van Leeuwen and Rabouille, 2019).

Here, we seek out to i) identify the pathways elicited in S2 cells by incubation in the starvation medium KRB that lead to Sec body formation, and ii) to assess whether the changes in the cytoplasmic biophysical properties play a role in the phase transition leading to Sec body formation. We show that amino-acid starvation in KRB stimulates the Unfolded Protein Response (UPR) and activates IRE1 autophosphorylation (Walter and Ron, 2011). However, the sole activation of the UPR does not lead to Sec body formation. To form Sec bodies, IRE1 phosphorylation needs to be combined to a moderate salt stress. In this regard, KRB incubation is faithfully mimicked by the cell incubation with DTT and addition of 100mM NaCl. Interestingly, a high salt stress (150mM NaCl) that activates the salt inducible kinases (SIKs) efficiently drives Sec body formation in a sufficient manner that is not due to osmotic shock and cell volume shrinkage. Importantly, we found that a 50% decrease in the ATP concentration is one factor strongly correlated to Sec body formation and partly depends on IRE1 kinase activity.

## Results

### Salt stress is necessary and sufficient for Sec body formation

In an attempt to understand Sec body formation that form upon incubation in the starvation buffer KRB (**Figure 1A,B**), we noticed that the salt concentration of the Schneider’s medium in which S2 cells are grown is much lower than mammalian tissue culture media, such as DMEM and KRB as used as a starvation medium (about 3-fold lower for Na^+^, **Table 1**). Accordingly, we found that lowering the concentration of NaCl in KRB (to 60mM instead of 120 in normal KRB) resulted in a decrease in Sec body formation (**Figure 1C**), showing that this salt is necessary for their formation. In this regard, the mere addition of 150mM NaCl together with 10mM sodium bicarbonate (SCH150, **Table 1**) to the Schneider’s leads to the solid formation of reversible Sec bodies, as efficiently as incubating the cells in KRB (**Figure 1A-C**).

**Table 1:** Ion concentrations in the different cell incubation media/buffers, ion concentration in KRB versus Schneider’s.

| Total | Schneider's (Sch) | KRB | SCH84 | SCH100 | SCH150 | KRB21 | DMEM |
| --- | --- | --- | --- | --- | --- | --- | --- |
| **Na<sup>+</sup> (mM)** | 51 | 138<br>(x2.7)* | 144<br>(x2.8) | 161<br>(x3.15) | 211<br>(x4.13) | 138<br>(x2.7) | 151<br>(x2.9) |
| **Cl<sup>-</sup> (mM)** | 67 | 126<br>(x1.9) | 151<br>(x2.3) | 168<br>(x2.5) | 218<br>(x3.3) | 142<br>(x2.1) | 126<br>(x2.2) |
| **K<sup>+</sup> (mM)** | 21 | 4.5<br>(x0.2) | 21<br>(x1) | 21<br>(x1) | 21<br>(x1) | 21<br>(x1) | 4.5<br>(x0.2) |
| **HCO<sub>3</sub><sup>-</sup> (mM)** | 5 | 15<br>(x3) | 15<br>(x3) | 15<br>(x3) | 15<br>(x3) | 15<br>(x3) | 29<br>(x6) |
| **Glucose (mM)** | 11 | 11 | 11 | 11 | 11 | 11 | 25 |
| **Amino-acids (mM)** | 78 | - | 78 | 78 | 78 | - | 10.4 |
| **Serum** | 10% | - | 10% | 10% | 10% | - | 10% |
| **Sec bodies** | No | Yes | No | No | Yes | Yes |  |

\*The (x) in bracket indicates the fold difference over the concentration in Schneider's.

Osmotic shock has been reported to trigger the formation of different stress assemblies, such as stress granules (Aulas et al., 2017) (Bounedjah et al., 2012) and P-bodies (Kilchert et al., 2010) (Jalihal et al., 2020). However, addition of 0.4-0.6M sucrose does not elicit Sec body formation (*Suppl Fig S1C, D*), even though the cell diameter significantly decreases by 16% (*Suppl Fig S1A, B*). This shows that the shrinkage of the cell volume is not a factor leading to Sec body formation. Furthermore, the addition of neither 150mM of KCl nor Na-acetate instead of NaCl leads to a solid formation of Sec bodies (*Suppl Fig S1C, D*). Taken together, these results suggest that the combined increase in the Na^+^ and Cl^−^ concentration in Schneider’s (4- and 3-fold, respectively, referred to, as salt stress)-but not osmotic shock or K^+^ (**Table 1**) triggers a pathway that leads to Sec body formation.

To test this further, we aimed to increase cytoplasmic Na^+^ concentration (in the presence of Cl^−^) and assess whether this is sufficient for Sec body formation. To do so, we focused on the abundant NaK ATPase present at the plasma membrane. NaK ATPase extrudes 3 Na^+^ against 2 K^+^, and is the main pump that maintains the intracellular Na^+^ concentration low and compatible with cellular function. We found that the localization of NaK ATPase changes dramatically from intracellular puncta in Schneider’s to strong plasma membrane localization in SCH150 (**Figure 1D**) together with a strong increase in its protein level (**Figure 1E**), suggesting its involvement. Accordingly, we found that incubation of cells with the NaK ATPase ouabain in growing medium SCH100 (**Table 1**) leads to a robust Sec body formation (**Figure 1 F, F’**), presumably because the cytoplasmic concentration of Na^+^ increases as it can no longer be extruded. This shows that increasing Na^+^ in the cytoplasm in the presence of Cl^−^ is sufficient to drive Sec body formation, perhaps by mobilizing the NaK ATPase.

We then asked which pathway is activated. An increase in intracellular Na^+^ concentration is known to activate the salt inducible kinases (SIKs). In this regard 3 SIKs are expressed in Drosophila (Teesalu et al., 2017) (Wehr et al., 2013) and we used a specific pan-inhibitor HG-9-91-01 (HG) to test their role in Sec body formation. Addition of HG reduces Sec body formation by 85% in cells incubated in SCH150 (**Figure 2A, A’**). Furthermore, a known target of SIK, HDAC4 (Histone Deacetylase 4) that is phosphorylated upon SIK activation is also activated in SCH150 (**Figure 2B**). Accordingly, addition of HG inhibits this phosphorylation. As HG can also inhibit P38 and Src (Clark et al., 2012), we used specific inhibitors to these two kinases and assessed their potential to inhibit Sec body formation upon SCH150. Neither the P38 inhibitor SB203580 nor the Src inhibitor Dasatinib, inhibit Sec body formation (**Figure 2A, A’**), showing that the effect of HG must be attributed to SIK inhibition.

**Figure 2:**
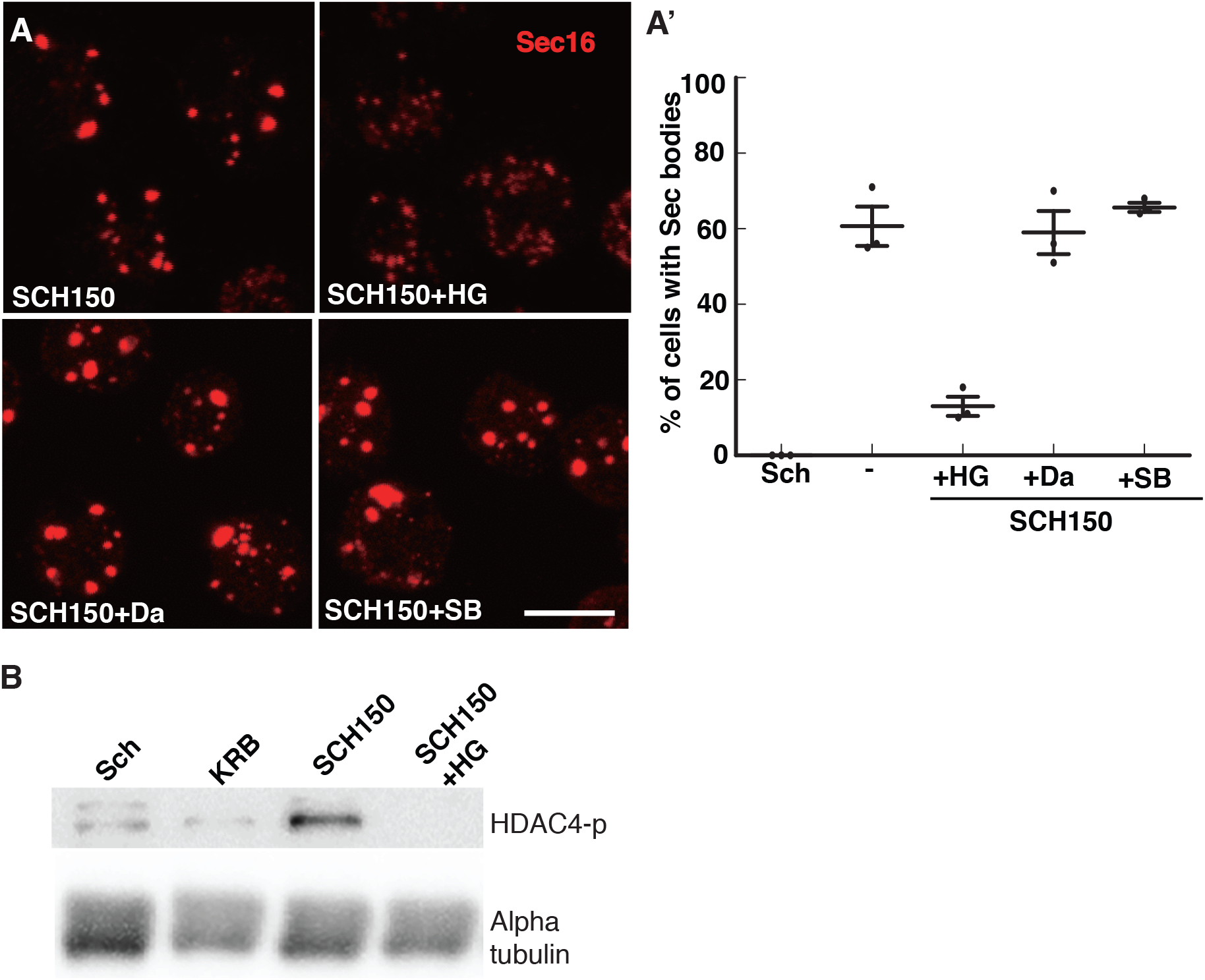
NaCl stress activates the salt inducible kinases that are involved in Sec body formation. **A, A’**: IF Visualization (A) and quantification (A’) of Sec body formation (marked by Sec16, red) in cells incubated in SCH150 supplemented or not with the SIK inhibitor HG-9-91-01 (HG, 5μM), the Src inhibitor dasatinib (Da, 20μM) and the p38MPKA inhibitor SB203580 (SB, 30μM) for 4h at 26°C. Note that the inhibition of Sec body formation is specific for HG-9-91-01. **B**: Western blot of S2 cells extract after incubation in Schneider’s (Sch), KRB and SCH150 with and without HG (5μM) for 4h at 26°C blotted for HDAC4-p and alpha tubulin. Note that the incubation with HG largely inhibits HADC-4 phosphorylation.

This not only shows that S2 cells are fully responsive to HG, but also that SIK activation is an important pathway for Sec body formation and that it is specifically activated by increased NaCl concentration. Taken together, this shows that the NaCl stress is a key pathway that leads to Sec body formation via SIK activation.

### In addition to salt stress, amino-acid starvation activates other pathways necessary for Sec body formation

The data above opens the possibility that the formation of Sec bodies in the starvation medium KRB is simply be due to an elevated salt concentration (**Table 1**), unrelated to the absence of amino-acids. However, this is not the case. When the Na+ and K+ concentration in Schneider’s and KRB are set to the same values (around 140 and 21mM, respectively, compare SCH84 and KRB21, **Table 1**), Sec bodies form very efficiently in KRB21 but not at all in SCH84 (**Figure 3A’**). The major difference between both is that SCH84 contains 78mM amino-acids and serum, whereas KRB contains none. Since we have demonstrated earlier that the serum (that was dialyzed to remove free amino-acids) does not play a role in Sec body formation (Zacharogianni et al., 2014) (Aguilera-Gomez et al., 2016), this indicates that the presence of amino-acids prevents Sec body formation even upon a moderate NaCl stress.

**Figure 3:**
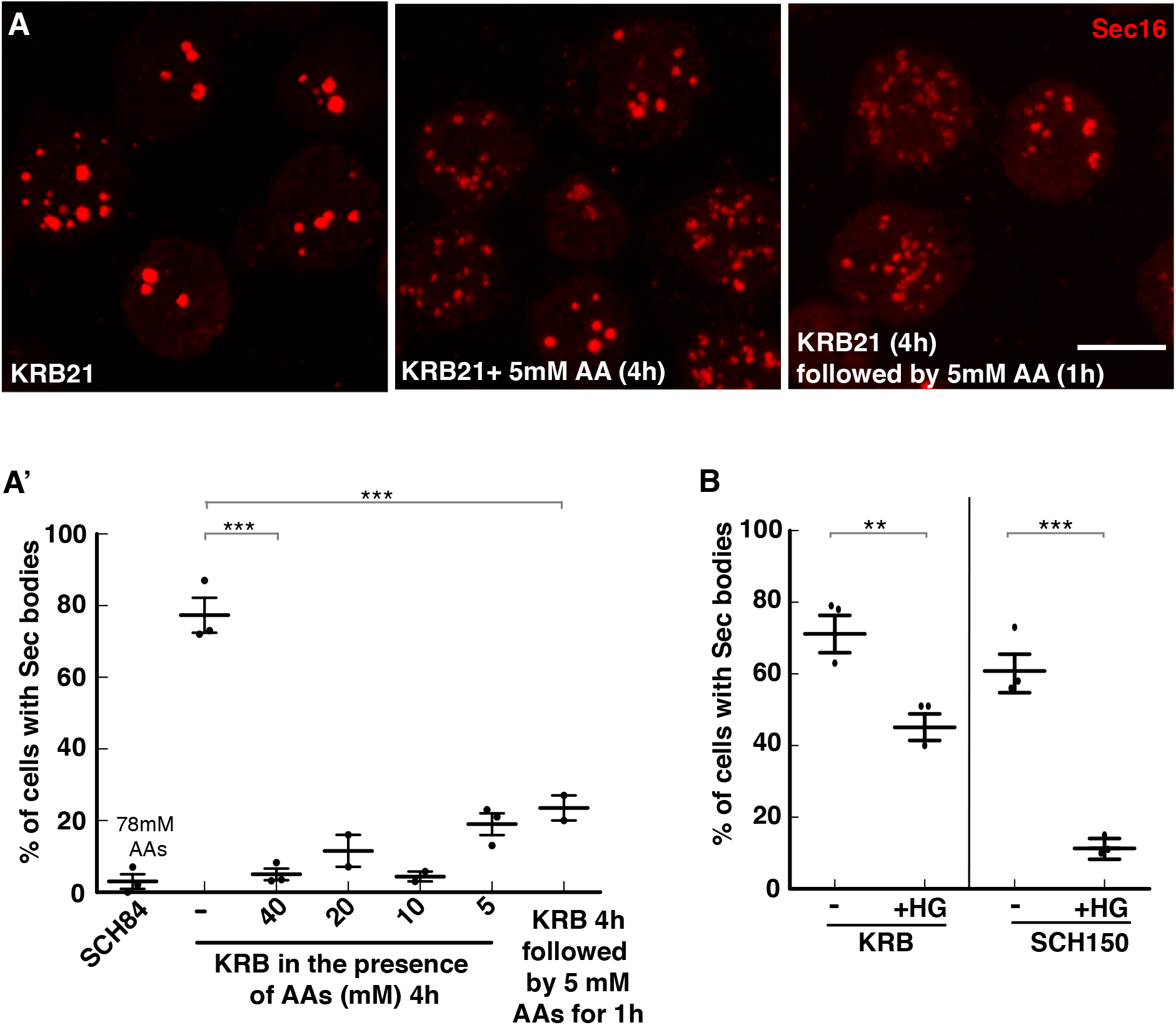
The presence of amino-acids prevents Sec body formation. **A, A’**: IF visualization (A) and quantification (A’) of Sec body formation (marked by Sec16) in cells incubated in SCH84, KRB21, KRB21 supplemented by 5-40mM amino-acids for 4h at 26°C, as well as KRB21 for 4h followed by addition of 5mM amino-acids for 1h at 26°C. Note that Sec bodies largely disappear when amino-acids are present. **B**: Quantification of Sec body formation (marked by Sec16) in cells incubated in KRB and SCH150 with or without the SIK inhibitor HG (5μM) for 4h at 26°C. Note that with the presence of HG inhibits 80% of Sec body formation in SCH150, but only 38% in KRB. Scale bar: 10μm Errors bars: SEM

To demonstrate this further, cells were incubated with KRB21 supplemented or not by amino- acids. Addition of as low as 5mM amino-acids to KRB21 strongly prevents Sec body formation (**Figure 3A, A’**), and revert KRB triggered Sec bodies back to ERES (**Figure 3A, A’**), almost as efficient as the reversion in Schneider’s (Zacharogianni et al., 2014). This suggests that for an equivalent salt concentration, the presence of amino-acids indeed prevents Sec body formation. This shows that amino-acid starvation is instrumental to Sec body formation.

Taken together, these results show that the absence of amino-acids in KRB either potentiates the salt stress (through increased Na^+^ and Cl^−^ concentration) or triggers another stress, leading to the formation of Sec bodies. This later hypothesis is strengthened by the fact that HG inhibits KRB triggered Sec body formation by 38%, not 80% as in SCH150 (**Figure 3B**). We therefore investigated which other pathways are triggered upon amino-acid starvation.

### The sole inhibition of mTORC1 is neither sufficient nor necessary to trigger Sec body formation

mTORC1 (mechanistic Target of Rapamycin complex 1) is the major sensor of amino-acid level in the circulating medium (Kim and Guan, 2019). When amino-acids are absent or low, the complex is inhibited resulting in the inhibition of many anabolic pathways, and the activation of the degradative pathway of autophagy. We have previously shown that in S2 cells incubated with KRB, protein synthesis is inhibited very quickly, the mTORC1 target S6 Kinase is no longer phosphorylated (*Suppl Figure S2A*), and the autophagic pathway is activated (Zacharogianni et al., 2014) (Zacharogianni et al., 2014). We also showed that the sole inhibition of mTORC1 by Rapamycin (Zacharogianni et al., 2014) (Zacharogianni et al., 2014) and Torin (*Suppl Figure S2B, B’*) is not enough to trigger the formation of Sec bodies. Similarly, depleting Raptor, the main subunit of mTORC1, does not trigger Sec body formation in growing cells (Zacharogianni et al., 2014) (Zacharogianni et al., 2014). Last, overexpression of TCS1/2, an endogenous inhibitor of mTORC1 (Condon and Sabatini, 2019) also failed to trigger the formation of Sec bodies (*Suppl Figure S2C*). Taken together, although amino-acid starvation does inhibit mTORC1 in S2 cells, Sec body formation is not a consequence of its sole inhibition.

### Amino-acid starvation leads to oxidative stress but oxidative stress alone is not enough to trigger Sec body formation

In order to figure out which other pathways are stimulated by amino-acid starvation in KRB, we performed bulk RNA sequencing and compared it to cells grown in Schneider’s. We detected 15688 genes, including 1022 downregulated and 945 upregulated (*Suppl Figure S3A, Suppl Table S1*, and GEO: GSE143810) [NCBI tracking system #20585561]). Interestingly, the strongest GO term associated to upregulated genes is oxidoreductase activity, including 20 glutathione-S-transferase as well as a strong reduction in peroxiredoxin expression, suggesting that KRB could elicit oxidative stress (*Suppl Figure S3B*), perhaps through ROS production.

To visualize the potential ROS production upon KRB incubation, we used DCF fluorescence measurements. KRB incubation elicits a robust and steady increase of ROS production that does not occur in Schneider’s. Yet, generating oxidative stress with either arsenite (*Suppl Figure S3* D, D’), APS or H_2_O_2_ (**not shown**) in growing cells does not lead to Sec body formation. Taken together, ROS are produced during KRB incubation but this is not sufficient to trigger Sec body formation.

### Amino-acid starvation stimulates the UPR via IRE1, but UPR is not enough to trigger Sec body formation

Since Sec bodies are linked to the inhibition of the exit of newly synthesized proteins out of the ER, we investigated whether incubation with the starvation medium stimulates the Unfolded Protein Response (UPR). We reasoned that this could be activated in response to an overload of misfolded proteins in the ER. UPR triggers the activation of three signaling pathways, the first and best characterized being the activation of IRE1. The misfolded protein overload leads to the dimerization of the IRE1 luminal domain resulting in the autophosphorylation of its kinase domain. This in turn, stimulates its nuclease activity. This leads to the specific splicing of *xbp1* mRNA that then translates into a potent transcription factor and that transcriptionally upregulates molecular machineries to resolve the overload of misfolded proteins in the ER (Walter and Ron, 2011). IRE1 nuclease activity is also required for RIDD (RNA IRE1 Dependent Degradation), a more unspecific degradation of mRNAs that are associated to the ER (Hollien et al., 2009).

Cells incubated in the starvation medium KRB displays a clear spliced *xbp1* (s, about 50% of total) when compared to growing cells where it is absent, showing that the IRE1 branch of the UPR is activated (**Figure 4A**). In line with it, IRE1 is clearly phosphorylated upon incubation in KRB (**Figure 4B**). However, RIDD did not appear to be stimulated by KRB incubation, at least when using *sparc* mRNA as a readout (Gaddam et al., 2012), (**Figure 4A**). In the absence of antibodies to Drosophila IRE1, we tested whether the expression level of the kinase was modified by KRB treatment by monitoring the level of its mRNA by PCR. However, we found that this treatment did not lead to an increase (**Figure 5D**).

**Figure 4:**
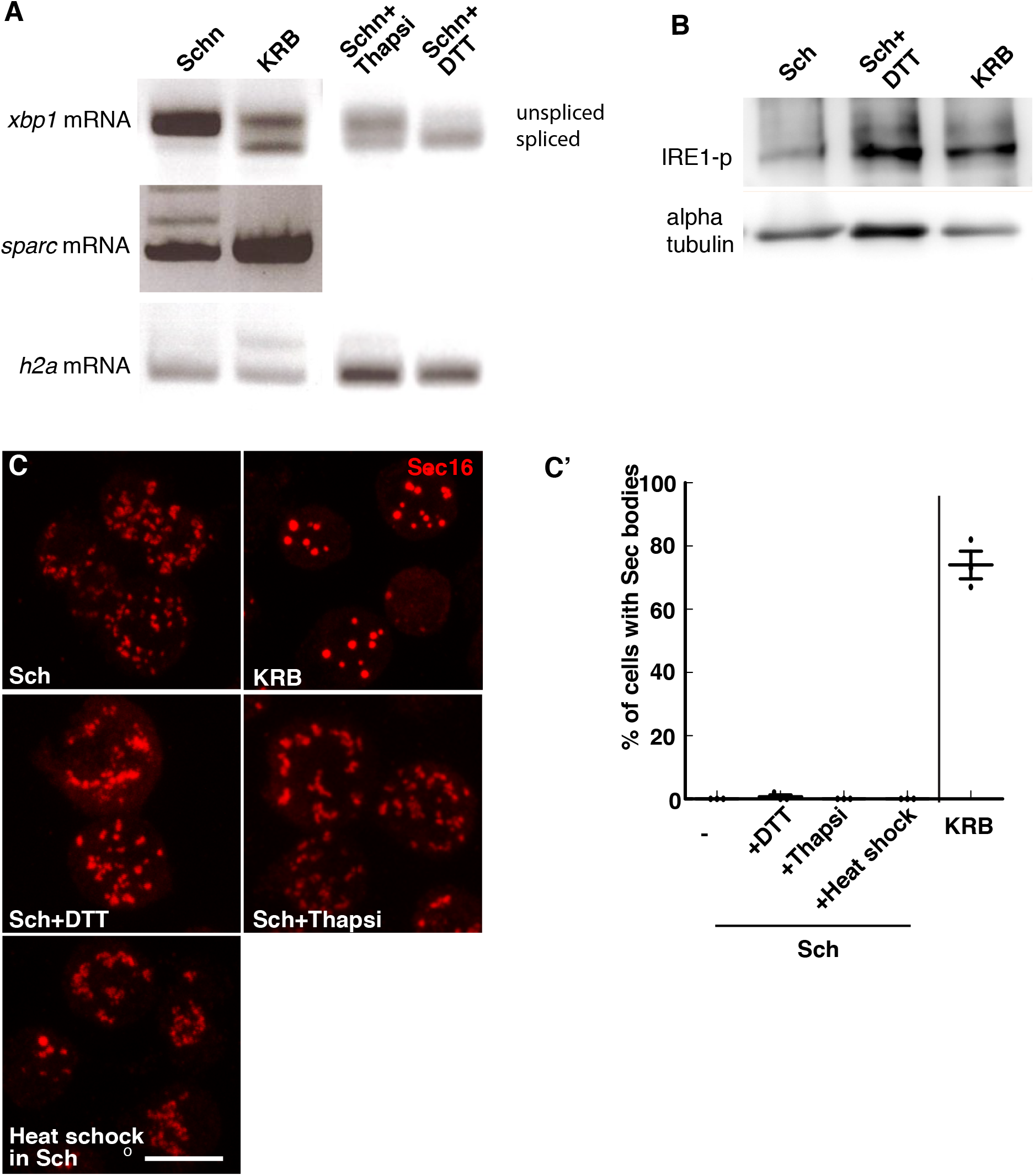
Amino-acid starvation in KRB activates the UPR and IRE1-phosphorylation. **A**: Visualization of the PCR products of *xbp1*, *sparc* and *h2a* mRNAs in cells grown in Schneider’s (Sch) and incubated in KRB for 4h. Note the robust *xbp1* splicing upon KRB. **B**: Western blot of S2 cells extract after incubation in Schneider’s, KRB and DTT (5mM) for 4h after blotting for IRE1-p (using an ab from Genetech). Note that KRB and DTT elicits IRE1 phosphorylation. **C, C’**: IF micrographs of Sec16 in cells in KRB, Schneider’s also supplemented with DTT (5mM), Thapsigargin (2μM) for 4h at 26°C, as well as incubated at 38°C (heat shock) for 4h. Note that, except for KRB, none of these conditions elicit Sec body formation. Quantification in C’. Scale bar: 10μm Errors bars: SEM

**Figure 5:**
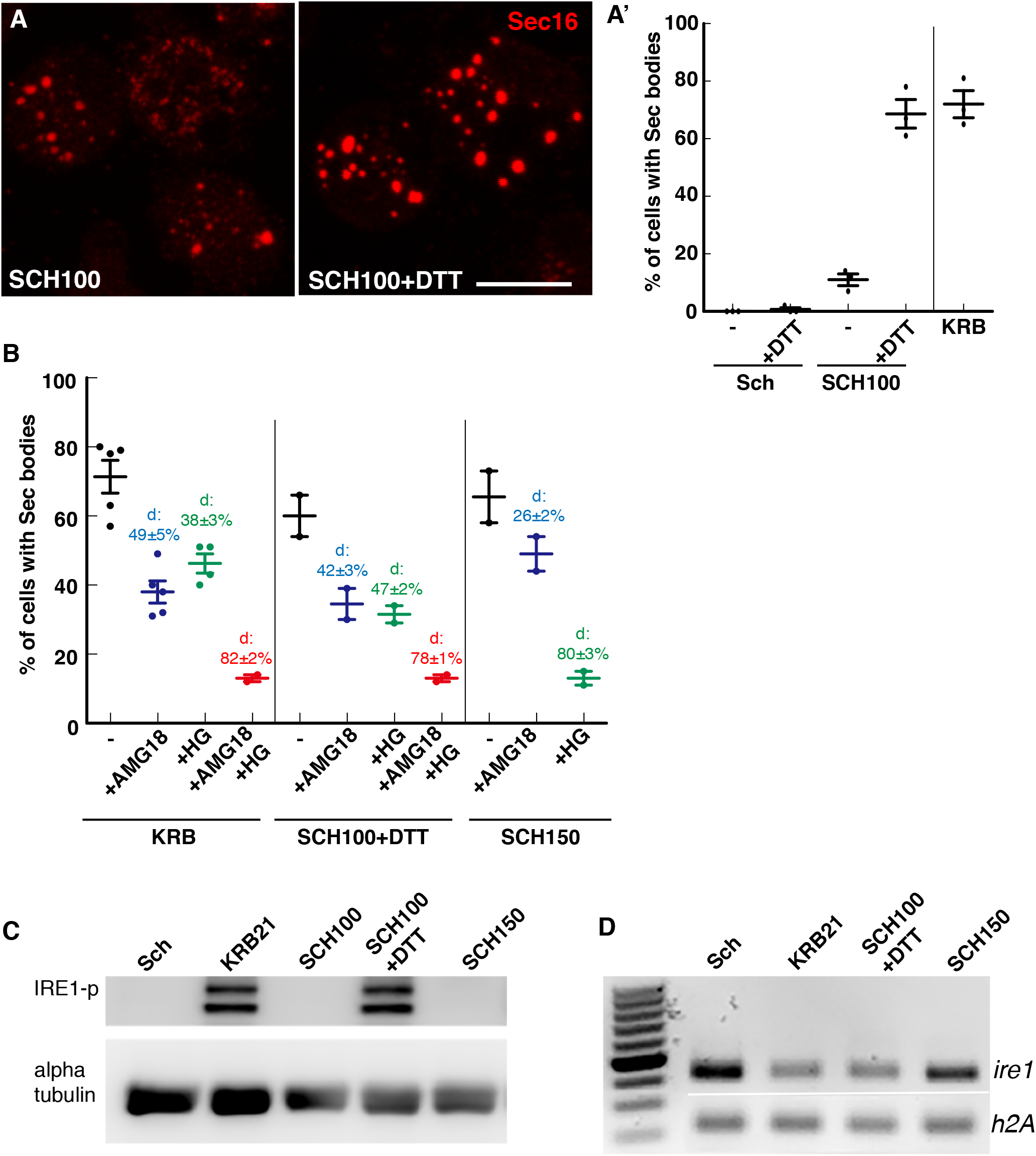
KRB incubation is mimicked by a moderate salt stress combined to ER stress. **A, A’**: Visualization (A) and quantification (A’) of Sec body formation (marked by Sec16) in cells incubated in KRB and SCH100+DTT (5mM) for 4h at 26°C. B: Quantification of Sec body formation (marked by Sec16) in cells incubated in the presence of the IRE1 kinase inhibitor AMG18 (10μM), the SIK inhibitor HG9-91-01 (HG, 5μM), and the combination of both upon KRB, SCH100+DTT and SCH150 treatment for 4h at 26°C. “d” indicates the decrease in Sec body formation observed upon treatment. P-value (SCH150 and SCH150+AMG18) is 0.1043. P-value (KRB and KRB+HG) is 0.0019. The other differences are highly significant (<10^−4^). **C**: Western blot of S2 cells extract after their incubation in Schneider’s, KRB and all the other conditions mentioned on the panel for 4h using the anti IRE1-p antibody (Genentech). **D**: Visualization of the PCR products of IRE1 and H2A mRNAs from cells incubated in Schneider’s (Sch), KRB21, SCH100+DTT (5mM) and SCH150 for 4h at 26°C. Scale bars: 10 μm Errors bars: SEM

Yet, activating the UPR alone through incubation with DTT and Thapsigargin (as exemplified by IRE1-p) (**Figure 4B**) and *xbp1* splicing (**Figure 4A**), and through heat shock does not lead to Sec body formation (**Figure 4C, C’**). Taken together, amino-acid starvation triggers the UPR and activates IRE1 but activation of this pathway is not sufficient to trigger Sec body formation, even when mTORC1 is also inhibited (**not shown**).

### A combination of moderate salt stress to ER stress leads to Sec body formation

As shown above, neither ROS production nor UPR/IRE1 activation on their own suffices to trigger Sec body formation. We then addressed whether combined with that NaCl stress, they would act in concert to activate Sec Body formation. We first tested a combination of moderate NaCl stress and ROS production, but this is largely not instrumental for Sec body formation (SCH100+Ars, *Suppl Figure S3D’*). The lack of involvement of ROS in Sec body formation was confirmed by inhibiting the ROS production in KRB with N-acetyl cysteine (NAC, *Suppl Figure S3C, D’*), which did not change the Sec body formation response.

We next tested whether the Sec body formation in KRB is recapitulated by applying a moderate NaCl stress together with activation of IRE1. Combining moderate salt stress (SCH100) with DTT (5mM, that induces ER stress and IRE1 auto-phosphorylation) (**Figure 5C**), is very efficient to induce the formation of largely reversible Sec bodies, to an extent that is similar to KRB (**Figure 5A**). Taken together, the Sec body formation in KRB is largely recapitulated by inducing a moderate salt stress together with UPR activation. The former stress would activate SIK, albeit partially (as in KRB), whereas the latter would lead to IRE1 autophosphorylation and activation.

To test further the comparison between SCH100+DTT and KRB, we inhibited IRE1 kinase activity with AMG18 (**Figure 5B**) combined or not with the SIK inhibitor HG. AMG18 incubation resulted in an about 45% decrease in the Sec body response in both conditions, showing that IRE1 kinase activity has the same contribution to Sec body formation both in KRB and SCH100+DTT. Furthermore, when IRE1 and SIK were inhibited together, the Sec body formation was strongly inhibited and in the similar manner in both KRB and SCH100+DTT (**Figure 5B**). This shows that moderate salt stress via SIK and ER stress via IRE1 combined are the two pathways driving Sec body formation. As a control, addition of AMG18 to SCH150 has only a moderate effect in agreement with the strong dependency on SIKs as shown by the strong inhibition by HG (**Figure 5B**). Of note, AMG18 is fully functional on Drosophila IRE1. First, the kinase site of Drosophila IRE1 is very similar to the human enzyme (*Suppl Figure S4A*). Furthermore, using AMG18 at either 10, 20 or 50μM during the KRB incubation inhibits Sec body formation to the same extent (*Suppl Figure S4B*). Last, the drug is 84% effective to block IRE1 phosphorylation by thapsigargin (*Suppl Figure S4C*). These results show that AMG18 is fully active in S2 cells.

Taken together, our results show that upon KRB incubation, Sec body form upon two pathways: The first is a moderate salt stress leading to the SIK activation. The second leads to IRE1 auto-phosphorylation and activation of its kinase domain due to the absence of amino-acids.

### The cytoplasm acidifies upon KRB incubation

As mentioned in the introduction, signaling pathways and/or change in the biophysical properties of the cytoplasm (Rabouille and Alberti, 2017) (van Leeuwen and Rabouille, 2019) are critical for phase separation leading to the formation of stress assemblies. We then questioned how the biophysical cytoplasmic features are affected by the activation of these two pathways.

In glucose starved yeast, the phase separation of proteasome subunits into Proteasome storage granules and the formation of metabolic enzymes foci have been shown to occur through a drop in the cytoplasmic pH in a necessary and sufficient manner (Peters et al., 2013) (Munder et al., 2016) (van Leeuwen and Rabouille, 2019). We then asked whether the cytoplasmic pH of the S2 cell cytoplasm also changed upon incubation in KRB and whether this is relevant to Sec body formation.

To do this, we used a cell-based assay to estimate the ratio of intensity of pHluorin versus mCherry in cells transfected with a pHluorin-mCherry cytoplasmic probe (Miesenböck et al.) (Brett et al., 2005). pHluorin is a pH-sensitive GFP mutant that changes its fluorescent spectrum upon a certain pH. Upon KRB incubation, we observed that the cytoplasmic pH decrease, illustrated by a decrease in the ratio pHluorin/mCherry when compared to cells grown in Schneider’s where it remains largely constant (*Suppl Figure S5A*, *A’*). We estimate that the cytoplasm of growing cells is at pH 6.8 whereas the pH of the cytoplasm of KRB incubated cells is 6 (*Suppl Figure S5A, A’*), suggesting a drop of nearly 1 pH unit. Interestingly, this result suggests that similar to yeast, nutrient starvation leads to a reduction of the cytoplasmic pH pointing to a conserved mechanism (Rabouille and Alberti, 2017) (van Leeuwen and Rabouille, 2019). However, we found that arsenite treatment of S2 cells also leads to a similar pH drop. However, this treatment does not lead to Sec body formation and this indicates that the decrease in pH is likely not a prime driver for Sec body formation.

### Sec body formation is largely associated to a drop in the cytoplasmic ATP concentration

In glucose starved yeast, the pH drop has been shown to be the consequence of a decrease in the intracellular ATP concentration (due to a reduction of glycolysis) (Munder et al., 2016). Using a luciferase-based assay, we therefore measured the intracellular ATP concentration in S2 cells that are amino-acid starved in KRB. The intracellular concentration is first stable for 1 hour, after which it steadily decreases, whereas it remains unchanged in cells kept in Schneider’s (**Figure 6A**). We estimate the intracellular ATP concentration to be 1.7mM (**Figure 6A, B,** *Suppl Figure S5B*) in growing cells, a concentration that decreases by 50% and 60 % after 3 and 4h incubation in KRB, respectively (**Figure 6A, B**). These results show that the sharp decrease in intracellular ATP concentration correlates to Sec body formation.

**Figure 6:**
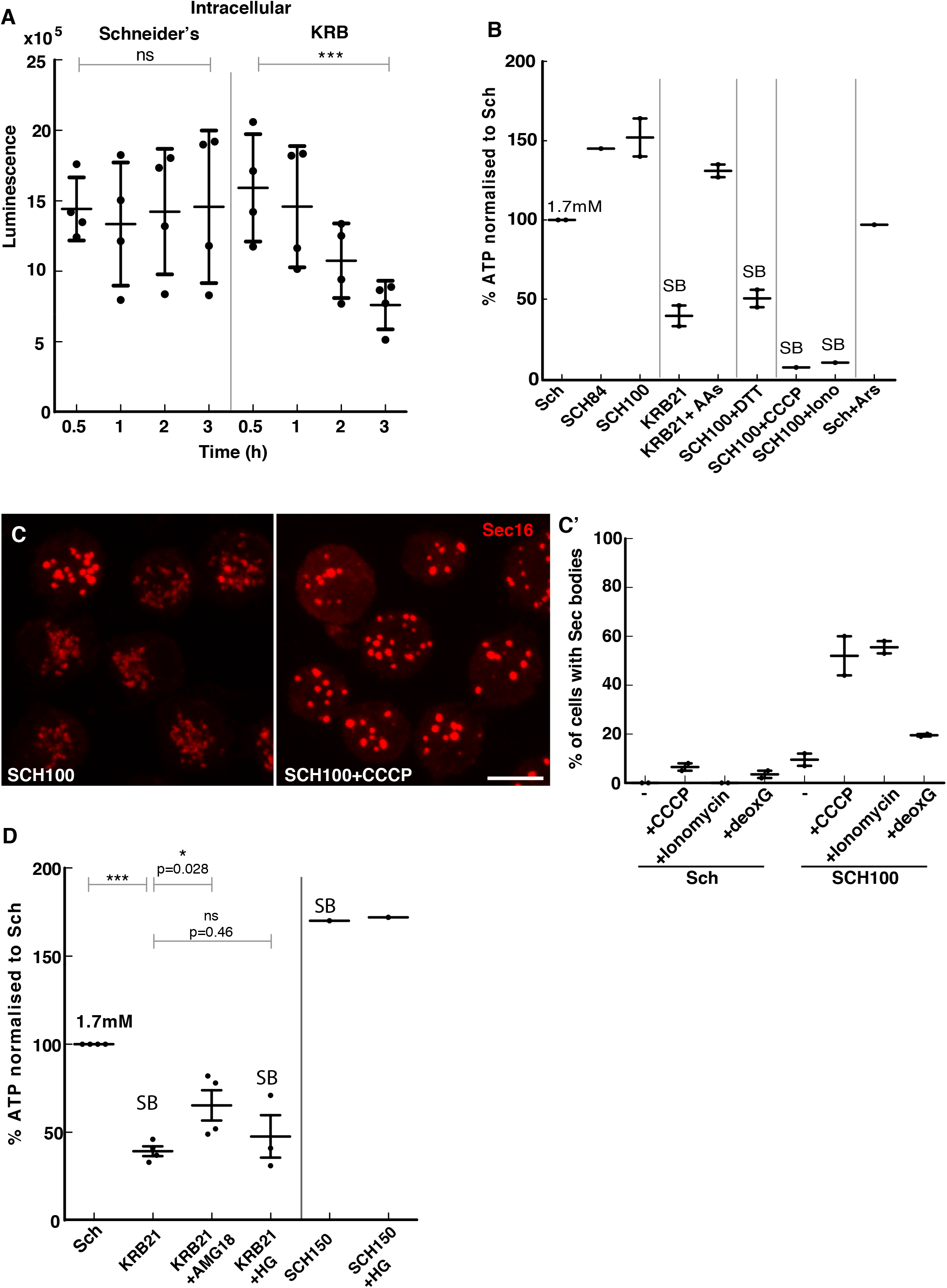
The decrease of the intracellular ATP level is one driving factor for Sec body formation. **A**: Luminescence intensity measuring the intracellular ATP concentration of S2 cells during incubation in Schneider’s (Sch) and KRB. Note that the ATP concentration decreases by 48% after 4h incubation in KRB. **B**: Quantification of the change in the intracellular ATP concentration (in % when compared to Sch that is set to 100%) after 4h incubation in the different conditions presented on the panel. **C, C’**: Visualization of Sec16 (C) and quantification (C’) of Sec body formation (marked by Sec16) in cells supplemented by CCCP (25μM), ionomycin (2.8μM) and 2-deoxyglucose (20mM) for 4h at 26°C. Note that inhibiting the OXPHOS leads to an efficient Sec body formation. **D**: Quantification of the change in the intracellular ATP concentration (in % when compared to Schneider’s that is set to 100%) upon IRE1-p inhibition with AMG18 (10μM) or SIK inhibition with HG-9-91-01 (5μM) in KRB21. The intracellular ATP was also measured upon SCH150 with and without SIK inhibition with HG-9-91-01 (5μM). Scale bar: 10μm Errors bars: SEM

To investigate further that the decrease in cytoplasmic ATP could be a driving factor in Sec body formation, we assessed the ATP level in experimental conditions that induce Sec body formation (**Figure 6B**). In particular, incubation in SCH100+DTT that leads to Sec body formation as KRB, also induces a severe drop in the intracellular ATP when compared to the control conditions where Sec bodies do not form. For instance, incubation with ouabain also leads to an ATP decrease (and Sec body formation, **Figure 1F, F’**). Conversely, in cells incubated in KRB supplemented with 5mM amino-acids where Sec body formation is largely inhibited when compared to KRB alone, the intracellular ATP concentration is not decreased (**Figure 6B**). Similarly, arsenite (that does not lead to Sec body formation) also does not lead to a drop in ATP. Overall, the conditions where Sec bodies form appears to be those where the ATP is the lowest.

To test whether a drop in intracellular ATP drives Sec body formation, we used two approaches. We first inhibited the ATP production of growing cells by blocking the mitochondrial OXPHOS by CCCP and ionomycin (**Figure 6C**) as well as by blocking glycolysis by 2-deoxyglucose. Strikingly, cells incubated in SCH100 supplemented by either CCCP or ionomycin leads to a robust Sec body formation, suggesting that the drop in intracellular ATP induces the formation of phase separated Sec bodies (**Figure 6C, C’**). Incubation with 2-deoxyglucose also induced a small and consistent Sec body formation (**Figure 6C’**), suggesting that S2 cells rely more on mitochondrial respiration than on glycolysis.

Second, we developed a semi-intact cell system using digitonin to gently permeabilize the S2 cell plasma membrane (**Figure 7A**). The specific incubation of the permeabilized cells with KRB at pH7.4 for 2h leads to Sec body formation in 38% of the cells (**Figure 7B, C**). We confirmed that the pH of the buffer and the presence of NaCl is critical, as KRB at pH6 and replacing Cl^−^ in the KRB with acetate did not lead to Sec body formation (**Figure 7B, C**). To test the role of a low concentration of ATP in Sec body formation, KRB was supplemented by 0.5mM ATP and this led to a strong inhibition of Sec body formation. This is specific for ATP as addition of AMP and Adenosine does not lead to this strong inhibition (**Figure 7D, D’**). These two results indicate that the intracellular ATP concentration modulates the phase separation of Sec bodies.

**Figure 7:**
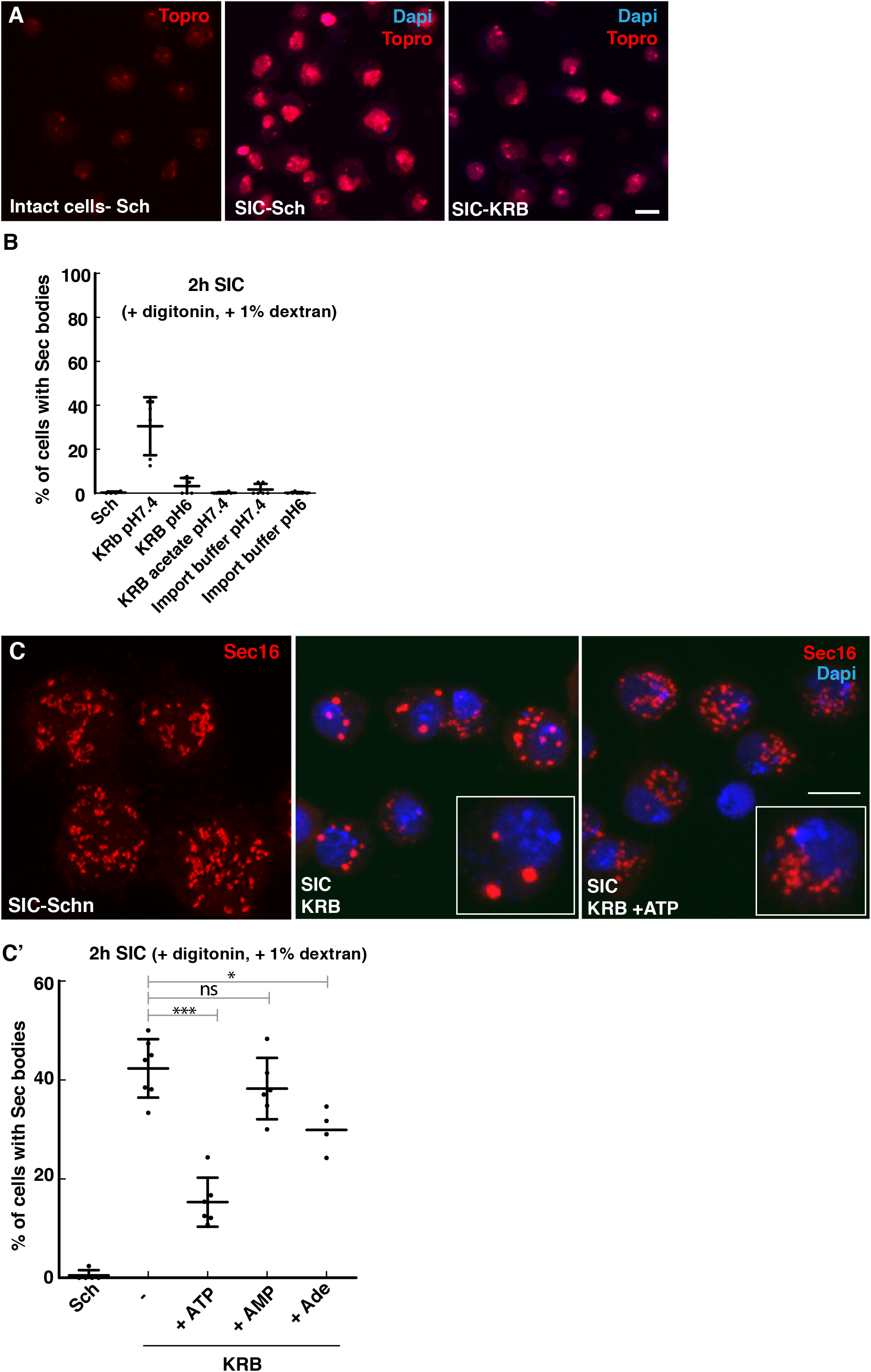
Addition of ATP to semi intact cells prevents Sec body formation. **A**: Visualization of S2 cells permeabilization using the non-membrane permeant TO-PRO-3 in intact cells in Schneider’s (Sch) and in semi-intact cells (SICs, i.e. + 10μg/ml digitonin, + 1% dextran) incubated in Sch and KRB for 2h at 26°C. Note that TO-PRO-3 stains the nucleus only in SICs. **B**: Effect of buffer composition in the formation of Sec bodies in SICs for 2h at 26°C. Decreasing the pH of KRB from 7.4 to 6 decreases the efficiency of Sec body formation. Replacing Cl^−^ by acetate in the KRB and replacing KRB by the import buffer (20mM HEPES, 110mM KAc, 2mM MgAc, 0.5mM EGTA) abolishes Sec body formation. **C, C’**: IF visualization of Sec16 (red, C) and quantification of Sec body formation (C’) (marked by Sec16) in SICs incubated in Sch, in KRB and in KRB supplemented by 0.5mM ATP, 0.5mM ADP and 0.5mM Adenosine. Note that Sec body formation is largely and specifically inhibited by addition of ATP. Scale bar: 10μm Errors bars: SEM (C), SD (B, C’).

Given the prominent role of IRE1 and SIK in Sec body formation, we tested whether the drop in intracellular ATP is linked to their activity. We first measured the ATP concentration in cells incubated in KRB in the presence of AMG18. The decrease in ATP concentration observed in KRB was less pronounced (**Figure 6D**) when IRE1 kinase activity was inhibited by AMG18, suggesting that this activity is involved in the observed ATP drop upon KRB incubation. This appears specific to IRE1 as addition of HG has only a small effect (**Figure 6D**), suggesting that SIK activity is not instrumental in this ATP drop. This is consistent with the noticeable fact that SCH150, a condition that leads to strong Sec body formation does not lead to a decrease in the cytoplasmic ATP concentration.

Taken together, these results strengthen the notion that the activation of two pathways are necessary for Sec body formation. The first is the SIK pathway that is sufficient (SCH150) and does not lead to a change in the cytoplasmic ATP concentration. The second is IRE1-p pathway that is necessary when the salt stress is moderate, but not sufficient. This pathway participates to the reduction of the cytoplasmic ATP, a factor we found is linked to Sec body formation

## Discussion

Here we show that Sec bodies in the Drosophila S2 cells form along two pathways (**Figure 8**).

**Figure 8:**
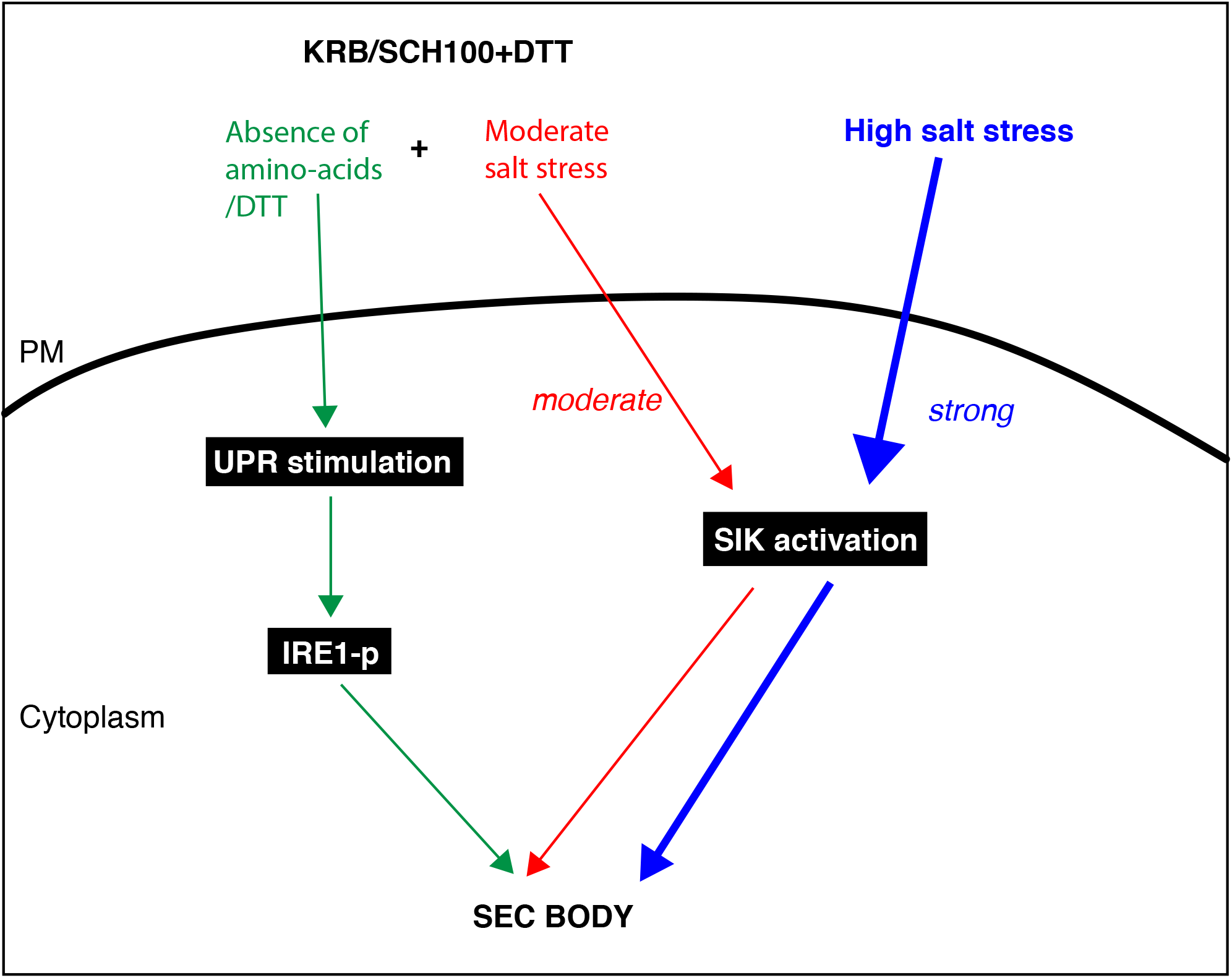
Activation of two pathways converging on IRE1 to form Sec bodies. Blue pathway: Sec bodies can form upon strong salt stress that activates the SIK in the cytoplasm. Red pathway: A moderate salt stress also activates SIKs, but it is not enough to stimulate Sec body formation on its own. Green pathway: Amino-acid starvation (or DTT) leads to IRE1-p that in turn, partly leads to ATP depletion. This needs to be combined with moderate SIK activation to lead to Sec body formation. Note that the blue pathway does not require ATP depletion for Sec body formation.

One is the activation of SIKs by high salt stress (addition to 150mM NaCl) that is necessary and sufficient. It is independent from osmotic shock and does not lead to a cell shrinkage. The other is a combination of a moderate salt stress and IRE1 activation (autophosphorylation triggered by the absence of amino-acids) that mimics the starvation buffer KRB. This is therefore well reproduced by addition of salt and DTT. Both leads to a 50% reduction in the cytoplasmic ATP concentration.

### The prominent role of Salt stress and SIK in remodeling the cytoplasm

Strong salt stress is induced by a 4-fold increase of Na^+^ in the medium combined with bicarbonate. This triggers an increase of Na^+^ in the cytoplasm that activates one or more salt inducible kinases (SIKs). SIK inhibition decreases Sec body formation.

Keeping intracellular Na^+^ as near as possible to physiological concentrations (5mM) is critical for cellular life, and the cell spends 40% of its available ATP to extrude Na^+^ against K^+^ through the NaK ATPase (Jaitovich and Bertorello, 2010) (our results, **Figure 6B**). It is therefore not surprising that sodium stress would elicit a cytoprotective response as prominent as Sec body formation (and stress granules in mammalian cells (Yan et al., 2014). This will need to be further elucidated as many organisms and tissues are subjected to increased circulating Na^+^. We show, however, that it is not equivalent to an osmotic shock and that this addition of salt does not lead to a decrease in a cell volume. Therefore, contrary to P-bodies (Kilchert et al., 2010) (Jalihal et al., 2020), this is not the mechanism by which Sec bodies are formed.

Increased Na^+^ activates the intracellular sodium-sensor network revolving around the SIKs (Jaitovich and Bertorello, 2010). The SIKs belong to the family of AMPK and have been shown to be part of a nutrient sensing mechanism so far revolving around glucose (Wein et al., 2018). In mammals, there are 3 genes encoding SIK (1-3) but only 2 in Drosophila: dSIK2 is the ortholog of human SIK1 and 2, and the 2 isoforms (long and short) of dSIK3 are homologs of human SIK3. dSIK2 has been shown to link to the fly Hippo pathway (Wehr et al., 2013), possibly linking nutrient to growth. dSIK3 is required for glucose sensing in the fly (Teesalu et al., 2017). Which SIK is involved in Sec body formation has not been clarified, as overexpression of each SIK individually has not proven enough to trigger Sec body formation, even when combined to some excess salt (SCH84 or 100). However, at least two of them appear to change their intracellular localization in KRB, i.e. SIK2 and long SIK3 isoforms, that appear to cluster near the plasma membrane and localize to the nuclear envelope (not shown). The role of SIKs in the formation of stress assemblies (here the Sec bodies) appears important and novel, and needs to be investigated further.

### IRE1 autophosphorylation

Although a high salt stress is sufficient to trigger Sec body formation, the Sec body formation observed during incubation in the amino-acid starvation buffer KRB elicits another pathway. Indeed, KRB induced Sec body formation is mimicked by a moderate salt stress (SCH100, **Table 1**) combined with the stimulation of the UPR through IRE1 activation by autophosphorylation by DTT (SCH100+DTT). Sec bodies that form in both conditions are similar in size and number and they are reversible. Whether their content is strictly similar has not been addressed here. Importantly, stimulation of the IRE1-p alone does not lead to Sec body formation.

IRE1 is an ER transmembrane sensor consisting of a luminal domain, a transmembrane domain and effector cytoplasmic portion that displays a kinase domain and RNase domain (Lee et al., 2008). What is now widely accepted is that upon ER stress driven by the overload of unfolded proteins, IRE1 luminal domain oligomerizes and this leads to the activation of IRE1 through autophosphorylation of its kinase domain (Ron and Walter, 2007). This activation is observed upon KRB incubation when the absence of amino-acids stimulates ER stress in a manner that is yet to be determined.

### Decrease in the intracellular ATP concentration is one driving factor in Sec body formation

We have shown that KRB leads to the decrease in the intracellular ATP concentration in Drosophila S2 cells. This is well reproduced by inhibition of the OXPHOS machinery of the mitochondria as well as using a semi-intact cell system. We show that this ATP decrease is an important factor, though not the sole one, for Sec body coalescence and we propose that it might be linked to its hydrotropic property, at least *in vitro*. Indeed, addition of 8mM ATP dissolves/prevents the phase separation of purified FUS and TAF15 (Patel et al., 2017), a concentration that is in the physiological range of ATP level in mammalian cells (between 1-10mM) (Traut, 1994) that is compatible with the S2 cell ATP concentration (1.7mM).

We found that this ATP concentration decrease is partly downstream of IRE1-p. How?

First, the ER and mitochondria are connected via the membrane contact sites coined “mitochondria-associated ER membrane” (van Vliet et al., 2014). Upon ER stress, Ca^2+^ is channeled from the ER to the mitochondria through these contacts and stimulate ATP production (Bravo et al., 2011). Excessive Ca^2+^ flux to the mitochondria (that can occur in ER stress upon KRB incubation) may lead to mitochondrial outer membrane permeabilization (Rizzuto et al., 2012), leading to the loss of these contact sites. As a result, the mitochondria would be disrupted and produces less ATP. Second, elevated Ca^2+^ levels in the mitochondria may also result in the activation of mitochondrial nitric oxide synthase (mtNOS), production of nitric oxide, S-nitrosylation and consequently to loss of mitochondrial function (Haynes et al., 2003) (Traaseth et al., 2004). One of the complexes that is inhibited by S-nitrosylation is Complex I of the electron transport chain leading in turn to a reduction in ATP levels (Morris et al., 2017). In this regard, it is interesting to note that nitric oxide exposure in mammalian cells leads to the formation of stress granules and a decrease in ATP levels (Aulas et al., 2018). Third, it is also relevant to mention that the unfolded protein response is a process highly demanding in ATP. Mostly the chaperones (e.g. BiP or calnexin) consume large amounts of ATP (Depaoli et al., 2019) (Vanhove et al., 2001). A fluctuation in ATP levels may prevent the chaperones from clearing misfolded proteins leading to an accumulation of misfolded proteins at the ERES.

Importantly, the sufficient high salt stress pathway that leads to Sec body formation does not lead to any decrease in the cytoplasmic ATP concentration, suggesting that, in addition to IRE1-p, another mechanism is activated that leads to a decrease in the cytoplasmic ATP. This remains to be identified.

In conclusion, this work illustrates the complexity of amino-acid starvation and the number of cellular parameters it affects. We propose that the formation of Sec bodies depends on the activation of signaling pathways leading to posttranslational modifications through activation of SIKs and IRE1 as well as the change in the cytoplasmic biophysical properties.

## Materials and Methods

### Cell culture, KRB incubation and drug treatments

Drosophila S2 cells were cultured in Schneider’s medium (Sch, Sigma) supplemented with 10% insect tested fetal bovine serum (Sigma) at 26°C. This medium is referred to as Schneider’s. S2 cells (between passages 5 and 18) were pelleted at 1200 rpm in a microfuge for 3 min, washed once in fresh Schneider’s, and diluted to 10^6^/ml. 1 ml of cell suspension were plated per well in a 12 wells plate containing coverslips. Cells were allowed to attach for 1.5h before starting the treatment.

Amino-acid starvation was performed in Krebs Ringers Bicarbonate buffer (KRB) comprising 0.7mM Na_2_HPO_4_, 1.5mM NaH_2_PO_4_, 15mM NaHCO_3_ (sodium bicarbonate, BIC), 120.7mM NaCl, 4.53mM KCl, 0.5mM MgCl and 10mM Glucose at pH7.4 as reported in (Zacharogianni et al., 2014) (**Table 1**). SCH84, SCH100 and SCH150 correspond to Sch supplemented to 84, 100 and 150 mM NaCl. KRB21 corresponds to KRB supplemented to 21mM KCl.

### Cell treatments

Drugs were used at the concentrations mentioned in *Suppl Table S2* including the IRE1 kinase inhibitor AMG18 (Amgen small molecule 18) (Harrington et al., 2015). The drug treatment was performed on the plated cells at 26°C for 4h incubated either in Schneider’s or in KRB or any of the modified versions of these two media (**Table 1**). Heat shock was performed on 2×10^6^ S2 cells plated on coverslip in 3 cm dish at 38 °C in an incubator for 1h.

### Immunofluorescence (IF)

Cells were fixed with 4% paraformaldehyde in PBS (pH 7.4) for 20 min. Cells were then washed 3 times with PBS and subsequently quenched by incubation in 50mM NH_4_Cl in PBS for 5 min. Followed by permeabilization with 0.11% Triton-X for 5 min. Hereafter, cells were washed 3 times in PBS and blocked in PBS supplemented with 0.5% fish skin gelatin (Sigma-Aldrich) for 20 min. Cells were then incubated with the primary antibody (in blocking buffer) for 25 min, washed 3 times with blocking buffer and incubated with the secondary antibody (in blocking buffer) coupled to a fluorescent dye for 20 min. Cells on the coverslip were washed 2 times in milliQ and dried for 3 min on a tissue with cells facing up. Finally, each coverslip with cells was mounted with Prolong antifade media (+DAPI, Thermofisher) on a microscope slide. Samples were viewed with a Leica SPE confocal microscope using a 63x oil lens and 2x zoom.

### Antibodies

For immunofluorescence, we used the rabbit polyclonal anti-Sec16 (1:800, (Ivan et al., 2008)) to detect Sec16 and the mouse monoclonal antibody a5 (1:20, DSHB) to detect NaK-ATPase. Donkey anti-rabbit Alexa 568 (1:200, Invitrogen) and a goat anti-mouse Alexa 488 (1:200, Invitrogen) were used as secondary antibodies.

For Western Blot, we used a rabbit monoclonal directed against the human IRE1-phosphorylation site (1:1000, gift from Genentech (Chang et al., 2018), a mouse monoclonal antibody a5(1:1000, DSHB) to detect NaK-ATPase, a rabbit monoclonal anti-phospho-HDAC4/HDAC5/HDAC7 (1:1000, Cell Signaling) and a mouse monoclonal anti-alpha-tubulin (1:2500, Sigma-Aldrich) followed by anti-rabbit and mouse antibodies coupled to HRP (1:2000, GE Healthcare).

### Quantification of Sec body formation

Cells positive for Sec bodies contain at least 2 large (>0.5μm) (typically 3-10) round Sec16 positive structures, whereas the typical ERES pattern observed in Schneider’s has largely disappeared (see **Figure 1A, A’**) (Zacharogianni et al., 2014). For all conditions, IF analysis was performed 3 times or more unless otherwise stated and at least 4 to 5 fields (about 25-30 cells per field) were recorded and analyzed per experiment. The response per field was determined by dividing the total amount of cells with Sec bodies by the total amount of cells. Finally, the average between all the fields was calculated and statistical analyses was performed using a one-tailed t-test.

### Cell diameter measurement

After incubation with different buffers, S2 cells were stained with Phalloidin-TRITC 1: 4000 (Sigma) for 20 min to detect F-actin. To measure the cell cross-sectional diameter, confocal images of equatorial cell sections were taken. The diameter was drawn by hand and measured using the “Measure” function of ImageJ.

### XBP1/RIDD assay and *IRE1* PCR

10×10^6^ cells grown in Schneider’s and 20×10^6^ cells incubated in KRB for 4h were spun down (5min 1400 rpm) and washed in PBS-MilliQ prior to RNA extraction. For each condition, RNA was extracted using the RNeasy Mini Kit (Qiagen). The RNA concentrations were measured and 1ug RNAs were used to synthesize cDNA using the GoScript Reverse Transcription System kit(Promega).

For each condition, a PCR was done using Taq polymerase (Promega) and visualize on agarose gel to assess *xbp1*, *sparc* (RIDD substrate), *IRE1* and *h2a* (control) cDNA.

Primers used are: XBP1 forward 5’-ATCAGCCAATCCAACGCCAG-3’, XBP1 reverse 5’-AGGCTCTTGCTGCTCTTCAG-3’, SPARC forward 5’-GTCGGACTGCTCTGTGTATC-3’, SPARC reverse 5’-ATGGTCCTTGTTGGAGTCGC-3’, H2A forward 5’-GTGGAAAAGGTGGCAAAGTGAA-3’, H2A reverse 5’-TTCTTGTTGTCACGAGCAGCAT-3’. IRE1 forward 5’-ATGAGAAGACGGACTGCACG-3’ IRE1 reverse 5’-GATCTGCTCGCCCTCCTTAC-3’.

### ROS detection assay

The stock solution of H2DCFDA (10mM in DMSO) was diluted to 5 μM in Schneider’s directly before usage. 1×10^6^ cells/ml were incubated with 5μM H2DCFDA in Schneider’s in the dark for 1h (an up to 4h) at 26°C in a 48-well plate. After incubation, the cells were washed 3 times with Schneider’s to remove excess H2DCFDA that might be noncovalent associated with the extracellular leaflet of the plasma membrane. The cells were then incubated in the treatment media (as described above) and the fluorescence intensity of the dye was immediately recorded over a period of 1h at 26°C using a Spark multimode microplate reader (Tecan) with an excitation of 480nm and an emission of 530nm. The fluorescence intensity of 5 fields of cells per condition was measured every 5 min for a period of 1h. Each experiment was performed at least 3 times or more.

The difference in DCF intensity for each condition was calculated by using the last value (1h) minus the first value (0 h). The difference in intensity of Schneider’s (usually very small, less than 10%) was used and set as the baseline at the value of 1. The difference in any other conditions was calculated as “Difference in DCF intensity _(condition x)_/ Difference in DCF intensity _(Schneider’s)_). The fold difference in was then expressed as a fold change over this baseline.

### Western Blot

A total of 4×10^6^ cells per condition were treated as described above. After treatment, the cells were harvested on ice and lysed in the following buffer (50 mM Tris-HCl pH 7.5, 150 mM NaCl, 1% Triton X-100, 50 mM NaF, 1 mM Na_3_VO_4_, 25 mM Na_2_-β-glycerophosphate supplemented with a protease inhibitor tablet (Roche). The lysates were cleared by centrifugation at 14,000 rpm for 20 min at 4°C. Supernatants were collected and the protein concentration was determined with a BCA protein assay kit (ThermoFisher). 50μg of protein was mixed with 5x SDS loading dye, boiled for 5 min and separated on an 8% SDS-PAGE gel. Then, separated proteins were transferred to a polyvinylidene difluoride (PVDF) membrane. Hereafter, the PVDF membrane was blocked in TBS+0.05%Tween-20 and 5% BSA (Sigma-Aldrich) (Blocking buffer). Primary antibodies were added to blocking buffer. After an overnight incubation at 4°C, the membrane was washed 3 times in TBS+0.05% Tween-20 over 45 min and incubated with secondary antibodies for 1 hour at room temperature. The membrane was washed 3 times washing in TBS+0.05%Tween-20 and developed by enhanced chemiluminescence (Bio-Rad) with Image Quant™ LAS 4000.

### Cytoplasmic pH measurement

The cytoplasmic pH was estimated by using the ratiometric fluorescent GFP-mutant pHluorin2 (Mahon, 2011) attached to mCherry in a cell-based assay. Cloning of pHluorin2-mCherry was performed by amplifying the fragment out of pAG304GPD-ypHluorin2 (Gift from Simon Alberti) using forward primer: 5’-GAGGGTACCATGGTGAGCAAGGGCG -3’ and reverse primer: 5’-CGACTCGAGTTATTTGTATAGTTCATCCATGCCA -3’ The fragment was cloned into the pMT-V5 vector using KpnI and XhoI restriction sites. pMT-pHluorin2-mCherry was transfected in S2 cells using Effectene transfection reagent (Qiagen) with a 1:10 ratio of DNA to Effectene Reagent.

Upon acidification, pHluorin2 excitation wavelength decreases, while the mCherry excitation signal remains largely unchanged. The ratio was calculated from the fluorescence intensity of pHluorin2 and mCherry determined using a Leica SP8 confocal microscope. The average ratio was calculated from 120 cells over 5 viewing fields.

A pH calibration curve was made by incubating transfected cells for 30 min in buffers with a different pH supplemented with 75μM monensin (Sigma-Aldrich), 10μM nigericin (Sigma-Aldrich), 10mM 2-deoxyglucose (Sigma-Aldrich) and 10mM NaN_3_ (Fisher scientific) to allow equilibration of the intracellular-extracellular-pH. The ratio was calculated as described above.

### ATP measurement

To measure the ATP concentration, we used the CellTiter-Glo 2.0 kit (Promega) and followed the manufacturer instruction. In brief, S2 cells (8×10^4^) were plated in a transparent 96-well plate and treated as indicated above. Every treatment was performed at least in triplicate and each experiment was performed 2-3 times. To determine the intracellular ATP concentration, 100ul of fresh incubation media was added to the treated cells and 0.5x volume CellTiter-Glo 2.0 (Promega) was added to each well. The background signal was determined by adding CellTiter-Glo 2.0 (Promega) to media without cells. The total volume was transferred to a 96-well Greiner flat white plate. Luminescence was measured with a Tecan Spark multimode microplate reader (emission wavelength 560nm).

To determine the extracellular ATP level, the incubation media was transferred to a 96-well Greiner flat white plate and 0.5x volume CellTiter-Glo 2.0 (Promega) was added to each well. The luminescence was measured as above. An ATP calibration curve was made by measuring different known concentrations of ATP (Sigma-Aldrich) in Schneider’s.

### Semi-intact cell assay (SIC)

WT S2 cells (1.5×10^6^) were plated on coverslips and permeabilized with 10 μg/ml digitonin (Sigma- Aldrich) in KRB for 2h at 26°C. Subsequently, cells were fixed and Sec bodies were visualized by immunostaining of Sec16. To test the effect of ATP, AMP or adenosine on Sec body formation, the semi-intact cells were incubated in the presence of 0.5mM ATP (Sigma-Aldrich), 0.5mM AMP (Sigma- Aldrich) or 0.5mM adenosine (Sigma-Aldrich). Note that the digitonin was not removed. The permeabilization efficiency was determined by using the non-membrane-permeable dye TO-PRO-3 iodide (ThermoFisher) (**Figure 7A**). The import buffer used in the semi-intact cell system comprised out of 20mM HEPES, 110mM KAc, 2mM MgAc, 5mM NaAc and 0.5mM EGTA (pH was set with KOH).

### RNA sequencing

A total of 10×10^6^ S2 cells were incubated in KRB and in Schneider’s for 4h as described above. Two biological replicates were performed. Total RNA was isolated using the RNeasy mini kit (Qiagen). The Utrecht Sequencing Facility (USEQ) generated RNA libraries using a RiboZero (Illumina) and TruSeq Stranded Total RNA Library Prep kit (Illumina). Hereafter, libraries were sequenced on Illumina NextSeq500 1 x 75 bp high output. Read quality was checked using fastqc. Reads were mapped to the Drosophila Melanogaster Dm6 reference genome (UCSC) using STAR (2.4.0.1) with default parameters (Dobin et al., 2013). We generated the read count per transcript table with an in-house Python2.7 script using pysam (https://github.com/pysam-developers/pysam) and the Dm6 annotation file (Ensembl). Differential expression between the conditions was determined using EdgeR (3.24.3) analysis in the R bioconductor package (Robinson et al., 2010). The DAVID web tool was used for GO- analysis with default parameters (https://david.ncifcrf.gov/summary.jsp). The accession number for the RNA-seq data reported in this paper is GEO (GSE143810) [**NCBI tracking system#20585561**].

### Statistics

Significance of the data was calculated and made by using GraphPad Prism 5.02. Statistical analyses were performed by using a one-tailed t-test. P-value is * when <0.05, ** when <0.01, *** when <0.001). Error bars represent SEM (except Fig 7B, C’ and Suppl Figure 5 where it represents SD).

## Supporting information

Suppl Materials Zhang et al 2021

## Acknowledgments

We thank our colleagues in the field for critics and comments. We thank Apfrida Kendek (UMCU) for Suppl Figure 2C. We acknowledge Genentech for providing us with the AMG18 and the anti IRE1-p monoclonal antibody.

---

*Suppl Figure S1: High salt stress is not equivalent to osmotic shock*

**A**: Visualization of the cell perimeter by Phalloidin staining (A) in cells incubated in Schneider’s, KRB and SCH150 for 4h at 26°C as well as Sch+0.4M sucrose for 1.5h at 26°C.

**B**: Measurement (in μm) of the cell diameter after incubations conditions mentioned on the panel. Note that incubation in 0.4M sucrose leads to a 16% decrease of the cell diameter.

**C**: IF visualization of Sec body formation (marked by Sec16) in cells in Schneider’s (Sch), SCH150, Sch supplemented with 10mM sodium bicarbonate and 150mM KCl, and with 0.4M sucrose. Note that except for SCH150, Sec body formation is inefficient.

**D**: Quantification of Sec body formation (marked by Sec16) in cells incubated in Schneider’s (Sch) and Schneider’s supplemented with 10mM sodium bicarbonate and 150mM of NaCl (SCH150), KCl, Na-acetate, and sucrose for 4h at 26°C. Note that the strong Sec body formation response is specific for addition of 150mM NaCl.

*Suppl Figure S2: TORC1 inhibition does not lead to Sec body formation*

**A**: Western blot of extract of S2 cells incubated as mentioned and blotted for S6K-p. This shows that S6K is no longer phosphorylated after KRB incubation for 4h and that rapamycin inhibits S6K-p stimulated by insulin.

**B, B’**: IF visualization (B) and quantification (B’) of Sec body formation (marked by Sec16, red) in cells grown in Schneider’s (Sch) also supplemented by mTORC1 inhibitors Rapamycin (10μM) and Torin (2μM) for 4h at 26°C Note that the sole inhibition of mTORC1 does not lead to Sec body formation contrary to incubation in KRB.

**C**: IF Visualization of Sec 16 in cells overexpressing the mTORC1 inhibitor TSC1/2-GFP. Note that Sec bodies do not form in these transfected cells.

Scale bar: 10μm

Error bars: SEM

*Suppl Figure S3: KRB incubation leads to oxidative stress*

**A**: Heatmap generated from all detected transcripts highlighting the similarities of the number of reads between the conditions based on the Euclidean distance. The replicates of each condition clusters together.

**B**: Volcano plot mRNAs identified by RNA sequencing that are differentially modulated between Schneider’s and KRB.

**C**: Graph showing the increase in DCF fluorescence measuring the production of ROS upon KRB21 for 4h at 26°C when compared to Schneider’s. Note that addition of N-acetyl-L-cysteine (NAC) suppresses ROS production.

**D, D’**: IF visualization (D) and quantification (D’) of Sec body formation in cells in Schneider’s supplemented with sodium arsenite (Ars, 2.5mM) and APS (500 μM) as well as SCH100+Ars (2.5mM) for 4h at 26°C. Note that none of these conditions elicit Sec body formation.

Scale bar: 10μm

Errors bars: SEM

*Suppl Figure S4: Drosophila IRE1 and specificity and efficacy of AMG18*

**A**: Comparison of the IRE1 catalytic sites sequences in human (H) and Drosophila (D). The green highlights the identity in and around the kinase site (underlined). The serine (S) in pink is the auto- phosphorylated site. Note that the two other serines (in blue) are also conserved.

**B**: Quantification of Sec body formation (marked by Sec16) in cells incubated in KRB supplemented by 10, 20 and 50uM AMG18. Note that the inhibition of Sec body formation is very similar for each concentration (about 45%).

**C**: Western blot of S2 cells extract after their incubation in Schneider’s supplemented with Thapsigargin (2μM) and AMG18 (10μM) using the IRE1-p specific antibody and an alpha tubulin antibody. Note that the incubation with AMG18 inhibits IRE1 phosphorylation by 84%

Errors bars: SEM

*Suppl Figure S5: pH measurements and calibration curves*

**A, A’**: Measurement of the intracellular pH (using the pHluorin-mCherry-fusion protein) in cells incubated in Schneider’s (Sch), KRB and Sch+0.5mM arsenite (A). pH calibration curve, plotting the ratio between pHluorin and mCherry in buffers with a different pH (A’)

**B**: ATP concentration calibration curve, plotting the ATP concentration versus the luminescence intensity.

Error bars: SD

*Suppl Table S1* (*related to Suppl Figure S3*): *List of genes up and down regulated upon KRB incubation.*

Differential gene expression between 4h starved (KRB) and 4h fed (Schneider’s) using EdgeR. The gene name, log fold change, P-value and FDR are shown. Significant (Empirical Bayes p<0.05) up regulated genes are highlighted in red and down regulated genes in blue.

*Suppl Table S2*: List of drugs with providers and concentrations

## Literature cited

Aguilera-Gomez, A., M.M. van Oorschot, T. Veenendaal, and C. Rabouille. 2016. In vivo vizualisation of mono-ADP-ribosylation by dPARP16 upon amino-acid starvation. eLife. 5.

Aulas, A., M.M. Fay, S.M. Lyons, C.A. Achorn, N. Kedersha, P. Anderson, and P. Ivanov. 2017. Stress-specific differences in assembly and composition of stress granules and related foci. Journal of cell science. 130:927–937.

Aulas, A., S.M. Lyons, M.M. Fay, P. Anderson, and P. Ivanov. 2018. Nitric oxide triggers the assembly of “type II” stress granules linked to decreased cell viability. Cell Death & Disease. 9:1129.

Bah, A., and J.D. Forman-Kay. Modulation of Intrinsically Disordered Protein Function by Post-translational Modifications.

Bounedjah, O., L. Hamon, P. Savarin, B. Desforges, P.A. Curmi, and D. Pastré. 2012. Macromolecular Crowding Regulates Assembly of mRNA Stress Granules after Osmotic Stress: NEW ROLE FOR COMPATIBLE OSMOLYTES. Journal of Biological Chemistry. 287:2446–2458.

Bravo, R., J.M. Vicencio, V. Parra, R. Troncoso, J.P. Munoz, M. Bui, C. Quiroga, A.E. Rodriguez, H.E. Verdejo, J. Ferreira, M. Iglewski, M. Chiong, T. Simmen, A. Zorzano, J.A. Hill, B.A. Rothermel, G. Szabadkai, and S. Lavandero. 2011. Increased ER–mitochondrial coupling promotes mitochondrial respiration and bioenergetics during early phases of ER stress. Journal of cell science. 124:2143.

Brett, C.L., D.N. Tukaye, S. Mukherjee, and R. Rao. 2005. The Yeast Endosomal Na+(K+)/H+ Exchanger Nhx1 Regulates Cellular pH to Control Vesicle Trafficking. Molecular biology of the cell. 16:1396–1405.

Chang, T.-K., D.A. Lawrence, M.M. Lu, J. Tan, J.M. Harnoss, S.A. Marsters, P. Liu, W. Sandoval, S.E. Martin, and A. Ashkenazi. 2018. Coordination between Two Branches of the Unfolded Protein Response Determines Apoptotic Cell Fate. Molecular cell. 71:629–636.e625.

Clark, K., K.F. MacKenzie, K. Petkevicius, Y. Kristariyanto, J. Zhang, H.G. Choi, M. Peggie, L. Plater, P.G. Pedrioli, E. McIver, N.S. Gray, J.S.C. Arthur, and P. Cohen. 2012. Phosphorylation of CRTC3 by the salt-inducible kinases controls the interconversion of classically activated and regulatory macrophages. Proceedings of the National Academy of Sciences of the United States of America. 42:1:6986–6991.

Condon, K.J., and D.M. Sabatini. 2019. Nutrient regulation of mTORC1 at a glance. Journal of cell science. 132:jcs222570.

Depaoli, M.R., J.C. Hay, W.F. Graier, and R. Malli. 2019. The enigmatic ATP supply of the endoplasmic reticulum. Biological Reviews. 94:610–628.

Dobin, A., D. C.A., F. Schlesinger, J. Drenkow, C. Zaleski, S. Jha, P. Batut, M. Chaisson, and T.R. Gingeras. 2013. STAR: ultrafast universal RNA-seq aligner. Bioinformatics. 29:15–21.

Farhan, H., M.W. Wendeler, S. Mitrovic, E. Fava, Y. Silberberg, R. Sharan, M. Zerial, and H.P. Hauri. 2010. MAPK signaling to the early secretory pathway revealed by kinase/phosphatase functional screening. The Journal of cell biology. 189:997–1011.

Gaddam, D., N. Stevens, and J. Hollien. 2012. Comparison of mRNA localization and regulation during endoplasmic reticulum stress in Drosophila cells. Molecular biology of the cell. 24:14–20.

Gomez-Navarro, N., and E. Miller. 2016. Protein sorting at the ER-Golgi interface. The Journal of cell biology. 215:769–778.

Harrington, P.E., K. Biswas, D. Malwitz, A.S. Tasker, C. Mohr, K.L. Andrews, K. Dellamaggiore, R. Kendall, H. Beckmann, P. Jaeckel, S. Materna-Reichelt, J.R. Allen, and J.R. Lipford. 2015. Unfolded Protein Response in Cancer: IRE1α Inhibition by Selective Kinase Ligands Does Not Impair Tumor Cell Viability. ACS Medicinal Chemistry Letters. 6:68–72.

Haynes, V., S. Elfering, R. Squires, N. Traaseth, J. Solien, A. Ettl, and C. Giulivi. 2003. Mitochondrial Nitric-oxide Synthase: Role in Pathophysiology. IUBMB life. 55:599–603.

Hollien, J., J.H. Lin, H. Li, N. Stevens, P. Walter, and J.S. Weissman. 2009. Regulated Ire1-dependent decay of messenger RNAs in mammalian cells. The Journal of cell biology. 186:323–331.

Ivan, V., G. de Voer, D. Xanthakis, K.M. Spoorendonk, V. Kondylis, and C. Rabouille. 2008. Drosophila Sec16 mediates the biogenesis of tER sites upstream of Sar1 through an arginine-rich motif. Molecular biology of the cell. 19:4352–4365.

Jaitovich, A., and A.M. Bertorello. 2010. Intracellular sodium sensing: SIK1 network, hormone action and high blood pressure. Biochimica et Biophysica Acta (BBA) - Molecular Basis of Disease. 1802:1140–1149.

Jalihal, A.P., S. Pitchiaya, L. Xiao, P. Bawa, X. Jiang, K. Bedi, A. Parolia, M. Cieslik, M. Ljungman, A.M. Chinnaiyan, and N.G. Walter. 2020. Multivalent Proteins Rapidly and Reversibly Phase-Separate upon Osmotic Cell Volume Change. Molecular cell. 79:978–990.e975.

Joo, J.H., B. Wang, E. Frankel, L. Ge, L. Xu, R. Iyengar, X. Li-Harms, C. Wright, T.I. Shaw, T. Lindsten, D.R. Green, J. Peng, L.M. Hendershot, F. Kilic, J.Y. Sze, A. Audhya, and M. Kundu. 2016. The Noncanonical Role of ULK/ATG1 in ER-to-Golgi Trafficking Is Essential for Cellular Homeostasis. Molecular cell. 62:982.

Kilchert, C., J. Weidner, C. Prescianotto-Baschong, and A. Spang. 2010. Defects in the secretory pathway and high Ca2+ induce multiple P-bodies. Molecular biology of the cell. 21:2624–2638.

Kim, J., and K.-L. Guan. 2019. mTOR as a central hub of nutrient signalling and cell growth. Nature cell biology. 21:63–71.

Lee, K.P.K., M. Dey, D. Neculai, C. Cao, T.E. Dever, and F. Sicheri. 2008. Structure of the dual enzyme Ire1 reveals the basis for catalysis and regulation in nonconventional RNA splicing. Cell. 132:89–100.

Mahon, M.J. 2011. pHluorin2: an enhanced, ratiometric, pH-sensitive green florescent protein. Adv Biosci Biotechnol. 2:132–137.

Miesenböck, G., D.A. D.A., and J.E. Rothman. Visualizing secretion and synaptic transmission with pH-sensitive green fluorescent proteins. Nature. 394:192–195.

Morris, G., M. Berk, H. Klein, K. Walder, P. Galecki, and M. Maes. 2017. Nitrosative Stress, Hypernitrosylation, and Autoimmune Responses to Nitrosylated Proteins: New Pathways in Neuroprogressive Disorders Including Depression and Chronic Fatigue Syndrome. Molecular Neurobiology. 54:4271–4291.

Munder, M.C., D. Midtvedt, T. Franzmann, E. Nuske, O. Otto, M. Herbig, E. Ulbricht, P. Muller, A. Taubenberger, S. Maharana, L. Malinovska, D. Richter, J. Guck, V. Zaburdaev, and S. Alberti. 2016. A pH-driven transition of the cytoplasm from a fluid- to a solid-like state promotes entry into dormancy. eLife. 5.

Owen, I., and F. Shewmaker. 2019. The Role of Post-Translational Modifications in the Phase Transitions of Intrinsically Disordered Proteins. International journal of molecular sciences. 20:5501.

Patel, A., L. Malinovska, S. Saha, J. Wang, S. Alberti, Y. Krishnan, and A.A. Hyman. 2017. ATP as a biological hydrotrope. Science. 356:753.

Peters, L.Z., R. Hazan, M. Breker, M. Schuldiner, and S. Ben-Aroya. 2013. Formation and dissociation of proteasome storage granules are regulated by cytosolic pH. The Journal of cell biology. 201:663–671.

Petrovska, I., E. Nuske, M.C. Munder, G. Kulasegaran, L. Malinovska, S. Kroschwald, D. Richter, K. Fahmy, K. Gibson, J.M. Verbavatz, and S. Alberti. 2014. Filament formation by metabolic enzymes is a specific adaptation to an advanced state of cellular starvation. eLife.

Rabouille, C., and S. Alberti. 2017. Cell adaptation upon stress: the emerging role of membrane-less compartments. Current opinion in cell biology. in press.

Rizzuto, R., D. De Stefani, A. Raffaello, and C. Mammucari. 2012. Mitochondria as sensors and regulators of calcium signalling. Nature Reviews Molecular Cell Biology. 13:566–578.

Robinson, M.D., M.D. J., and G.K. Smyth. 2010. edgeR: a Bioconductor package for differential expression analysis of digital gene expression data. Bioinformatics. 26:139–140.

Ron, D., and P. Walter. 2007. Signal integration in the endoplasmic reticulum unfolded protein response. Nature reviews. Molecular cell biology. 8:519–529.

Sprangers, J., and C. Rabouille. 2015. SEC16 in COPII coat dynamics at ER exit sites. Biochemical Society transactions. 43:97–103.

Teesalu, M., B.M. Rovenko, and V. Hietakangas. 2017. Salt-Inducible Kinase 3 Provides Sugar Tolerance by Regulating NADPH/NADP+ Redox Balance. Current Biology. 27:458–464.

Traaseth, N., S. Elfering, J. Solien, V. Haynes, and C. Giulivi. 2004. Role of calcium signaling in the activation of mitochondrial nitric oxide synthase and citric acid cycle. Biochimica et Biophysica Acta (BBA) – Bioenergetics. 1658:64–71.

Traut, T.W. 1994. Physiological concentrations of purines and pyrimidines. Mol Cell Biochem. 140:1–22.

van Leeuwen, W., and C. Rabouille. 2019. Cellular stress leads to the formation of membraneless stress assemblies in eukaryotic cells. Traffic. 0.

van Vliet, A.R., T. Verfaillie, and P. Agostinis. 2014. New functions of mitochondria associated membranes in cellular signaling. Biochimica et Biophysica Acta (BBA) - Molecular Cell Research. 1843:2253–2262.

Vanhove, M., Y.-K. Usherwood, and L.M. Hendershot. 2001. Unassembled Ig Heavy Chains Do Not Cycle from BiP In Vivo but Require Light Chains to Trigger Their Release. Immunity. 15:105–114.

Walter, P., and D. Ron. 2011. The Unfolded Protein Response: From Stress Pathway to Homeostatic Regulation. Science. 334:1081.

Wehr, M.C., M.V. Holder, I. Gailite, R.E. Saunders, T.M. Maile, E. Ciirdaeva, R. Instrell, M. Jiang, M. Howell, M.J. Rossner, and N. Tapon. 2013. Salt-inducible kinases regulate growth through the Hippo signalling pathway in Drosophila. Nature cell biology. 15:61–71.

Wein, M.N., M. Foretz, D.E. Fisher, R.J. Xavier, and H.M. Kronenberg. 2018. Salt-Inducible Kinases: Physiology, Regulation by cAMP, and Therapeutic Potential. Trends in Endocrinology & Metabolism. 29:723–735.

Wilhelmi, I., R. Kanski, A. Neumann, O. Herdt, F. Hoff, R. Jacob, M. Preussner, and F. Heyd. 2016. Sec16 alternative splicing dynamically controls COPII transport efficiency. Nature communications. 7:12347.

Yan, C., Z. Yan, Y. Wang, X. Yan, and Y. Han. 2014. Tudor-SN, a component of stress granules, regulates growth under salt stress by modulating GA20ox3 mRNA levels in Arabidopsis. Journal of Experimental Botany. 65:5933–5944.

Zacharogianni, M., A. Aguilera-Gomez, T. Veenendaal, J. Smout, and C. Rabouille. 2014. A stress assembly that confers cell viability by preserving ERES components during amino-acid starvation. eLife. 3.

