## Supplementary material for "Salt Inducible Kinase activation and IRE1-dependent intracellular ATP depletion to form Sec bodies in Drosophila cells": Suppl Materials Zhang et al 2021

*Suppl Figure S1. Zhang et al, 2021*

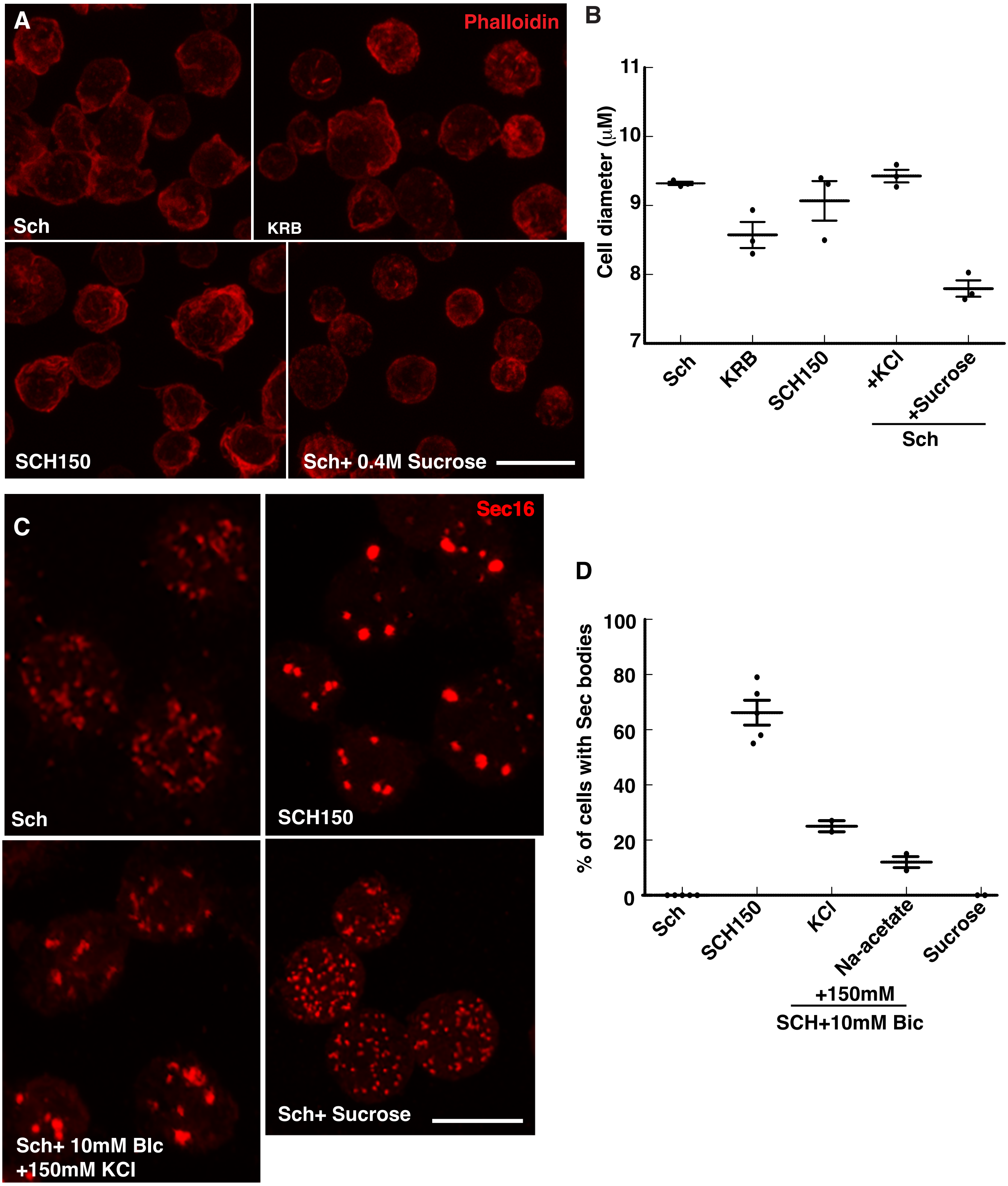

*Suppl Figure S2. Zhang et al, 2021*

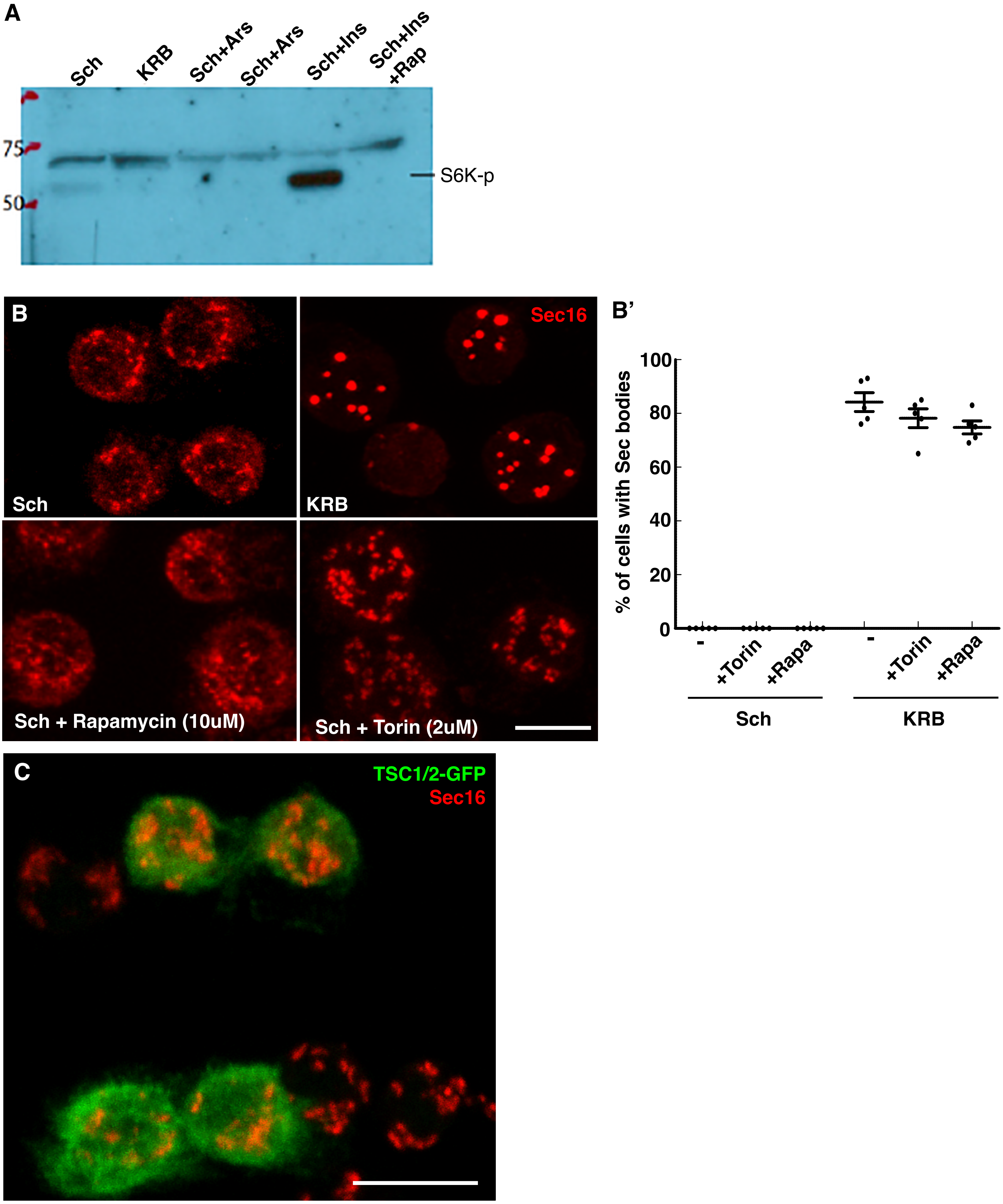

*Suppl Figure S3. Zhang et al, 2021.*

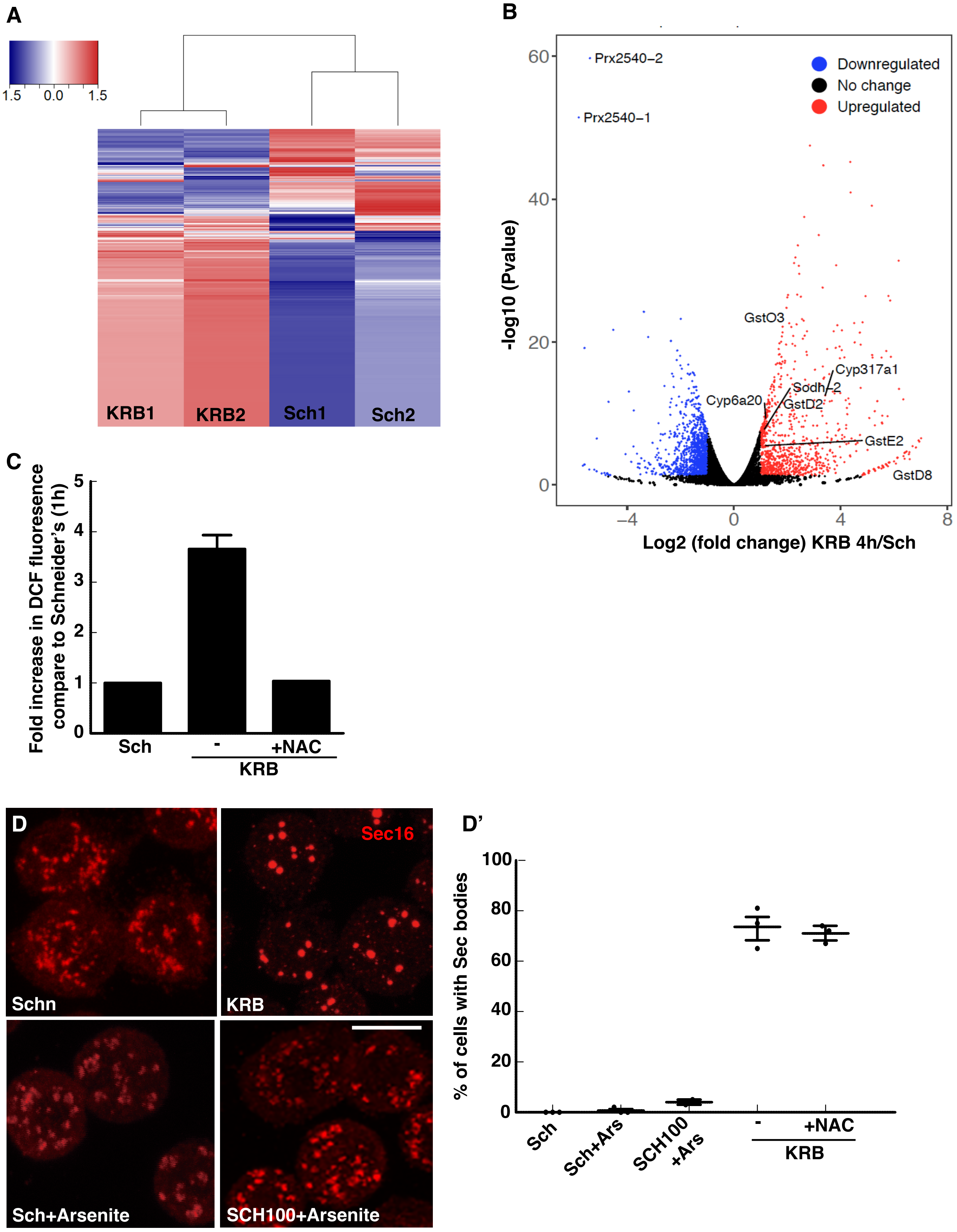

*Suppl Figure S3: Zhang et al, 2021.*

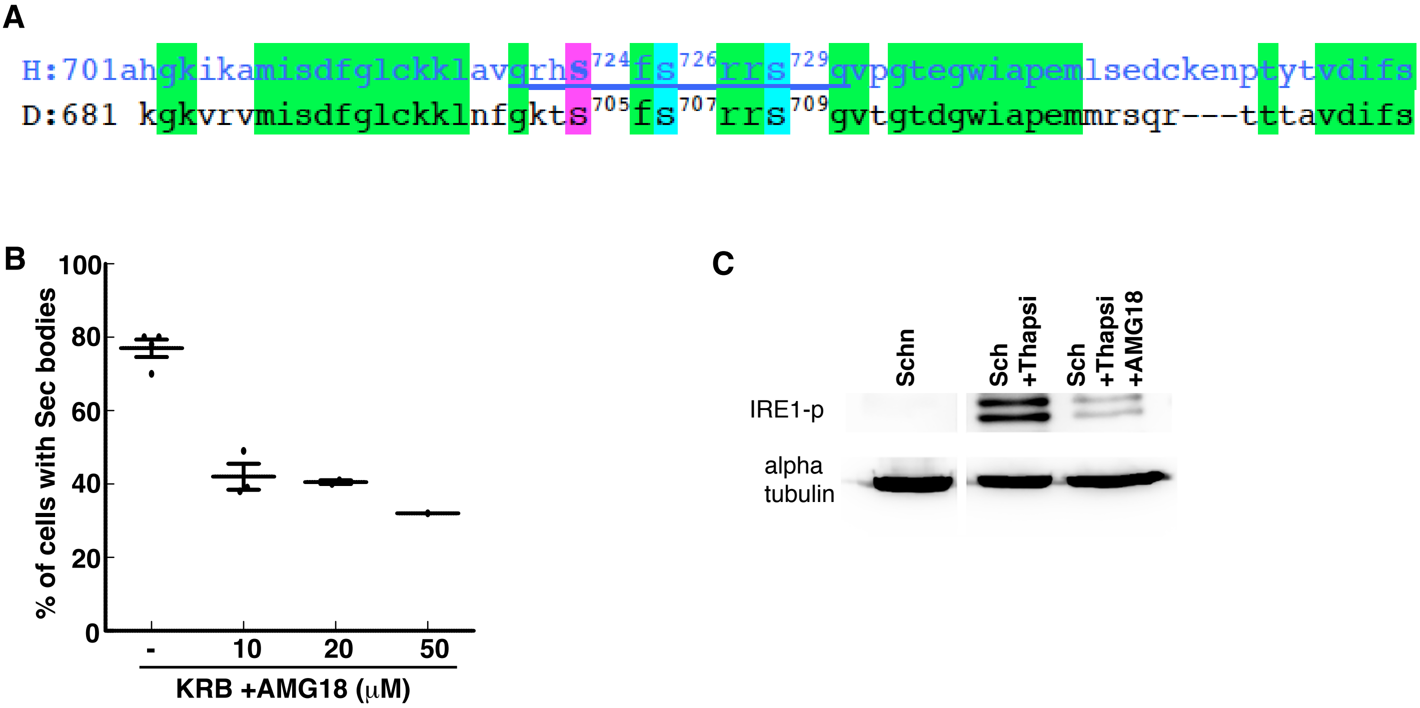

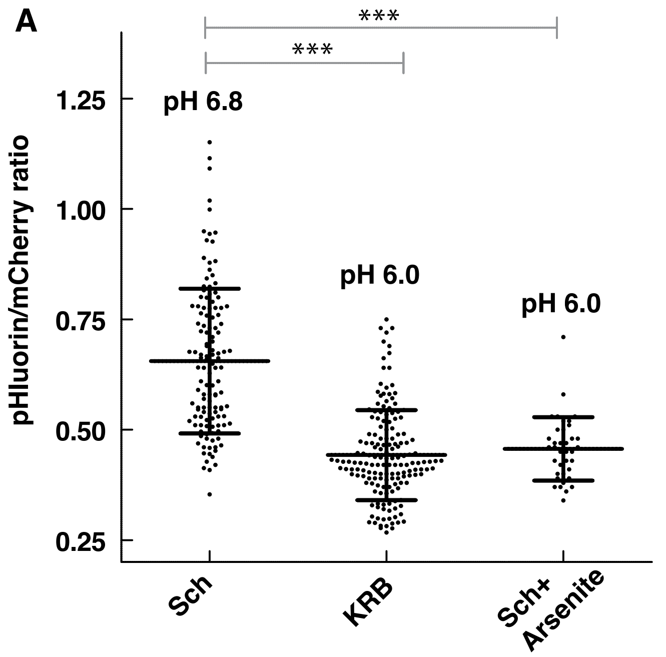
A
***
***
pH 6.8
pH 6.0
pH 6.0
pHluorin/mCherry ratio
1.25
1.00
0.75
0.50
0.25
Sch
KRB
Sch+ Arsenite

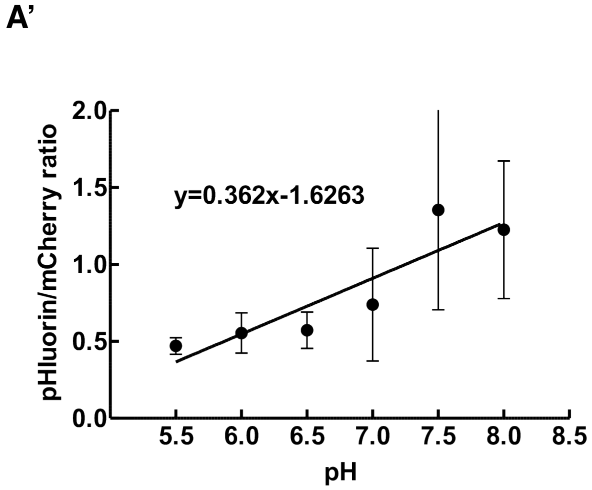
A'
pHluorin/mCherry ratio
y=0.362x-1.6263
2.0
1.5
1.0
0.5
0.0
5.5
6.0
6.5
7.0
7.5
8.0
8.5
pH

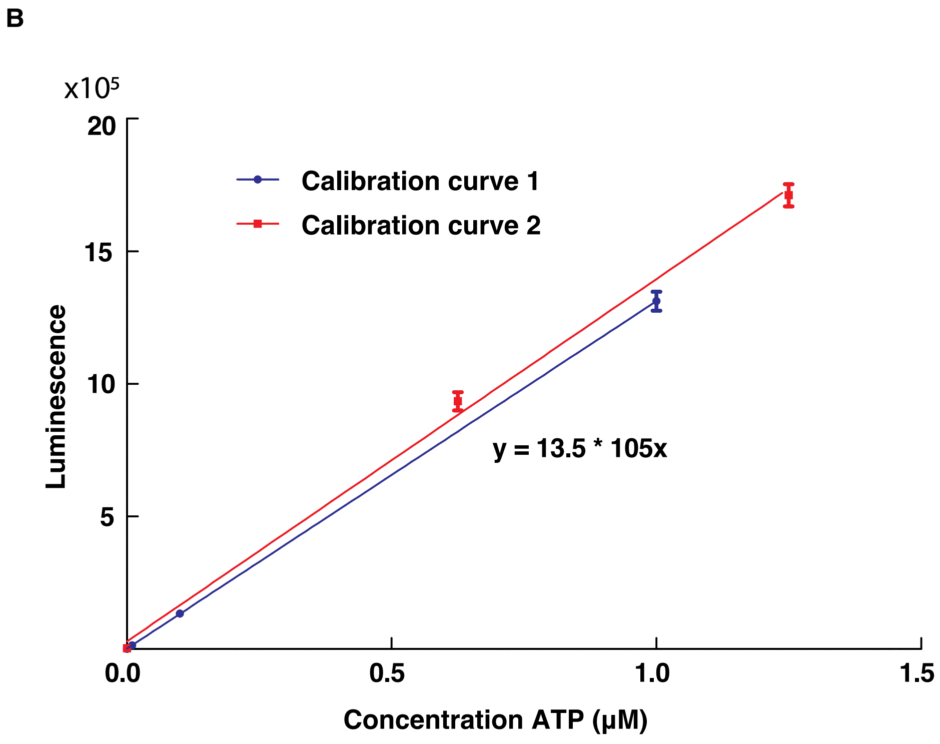
B
x10⁵
20
15
10
5
Calibration curve 1
Calibration curve 2
y = 13.5 * 105x
Luminescence
0.0
0.5
1.0
1.5
Concentration ATP (μM)

*Suppl Table S2: List of drugs with providers and concentrations.*

| **Chemicals** | **Suppliers** | **Stock concentrations** | **Solvant** | **Final concentrations** |
|---|---|---|---|---|
| Adenosine | Sigma-Aldrich | 10mM | MilliQ | 0.5mM |
| Amino-acids solution | Sigma-Aldrich | 228.7mM | KRB | 5.72mM |
| AMP (Adenosine 5'-monophosphate disodium salt) | Sigma-Aldrich | 100mM | MilliQ | 0.5mM |
| AMG18 (IRE1 kinase activity) | Gift from Genentech | 10mM | DMSO | 10µM<br>20µM<br>50µM |
| APS (Ammonium persulfate) | Sigma-Aldrich | 1M | MilliQ | 500µM |
| Arsenite (Na (meta) arsenite) | Sigma-Aldrich | 0.5M | MilliQ | 2.5mM |
| ATP (Adenosine 5'-triphosphate disodium salt hydrate) | Sigma-Aldrich | 100mM | MilliQ | 0.5mM |
| CCCP (Carbonyl cyanide 3-chlorophenylhydrazone) | Sigma-Aldrich | 100mM | DMSO | 25µM |
| Dasatinib | Sigma-Aldrich | 80mM | DMSO | 20µM |
| 2-Deoxy-D-glucose | Sigma-Aldrich | 1M | MilliQ | 20mM |
| DTT (Dithiothreitol) | Biorad | 2M | DMSO | 5mM |
| $H_2O_2$ (Hydrogen peroxide) | Sigma-Aldrich | 2M | MilliQ | 1mM |
| HG-9-91-01 (pan SIK inhibitor) | MedChemExpress/Bio-Connect | 5mM | DMSO | 5µM |
| KCl (Potassium Chloride) | Sigma-Aldrich | 2M | MilliQ | 150mM |
| Ionomycin from Streptomyces conglobatus | Sigma-Aldrich | 2.8mM | DMSO | 2.8µM |
| Na-acetate (Sodium acetate) | Fisher Scientific | 2M | MilliQ | 150mM |
| N-Acetyl-L-cysteine | Sigma-Aldrich | 30mM | MilliQ | 300µM |
| NaCl (Sodium Chloride) | J.T.Baker | 2M | MilliQ | 84mM,<br>100mM,<br>150mM |
| Ouabain octahydrate | Sigma-Aldrich | 50mM | MilliQ | 1µM |
| Rapamycin | Sigma-Aldrich | 10mM | DMSO | 10µM |
| SB203580 | Sigma-Aldrich | 30mM | DMSO | 30µM |
| Sucrose | Sigma-Aldrich | 2M | MilliQ | 0.4M |
| Thapsigargin | Sigma-Aldrich | 1mM | DMSO | 2µM |
| Torin | Invivogen | 3mM | DMSO | 2µM |
